# Cell-specific isotope labeling identifies *myo*-inositol transfer between neurons and oligodendroglia to support myelin repair

**DOI:** 10.64898/2026.03.19.712965

**Authors:** Kayla Adkins-Travis, Mun-Gu Song, Michaela Schwaiger-Haber, Kevin Cho, Ronald Fowle-Grider, Stephen L. Johnson, Leah P. Shriver, Gary J. Patti

## Abstract

Neurons and glial cells are biochemically coupled through the exchange of nutrients, but our knowledge of which metabolites are transferred between them remains limited due to technical challenges. Here, we introduce a strategy to label specific cell types with isotopic tracers so that metabolite transfer can be measured directly in the intact brain. By engineering neurons in mice to metabolize ^13^C-labeled cellobiose, a glucose dimer that wild-type cells cannot catabolize, we selectively track neuron-derived metabolites by using mass spectrometry-based metabolomics. Applying this approach enabled us to identify *myo*-inositol as a critical metabolite synthesized by neurons and transferred to oligodendrocyte progenitor cells (OPCs) via the SLC5A3 transporter. The transfer of *myo*-inositol from neurons to OPCs promotes OPC proliferation and differentiation by enhancing phosphatidylinositol synthesis and upregulating expression of myelin-associated genes. During demyelination, deficient nutrient transfer can be rescued by dietary supplementation of *myo*-inositol, which accelerates myelin repair. These findings establish a generalizable technology for tracing intercellular metabolite transfer *in vivo* and identify a previously unrecognized mechanism of *myo*-inositol transfer from neurons to glial cells in support of CNS regeneration, revealing a potential metabolic target for therapeutic intervention in neurodegenerative disease.

## Introduction

The brain lacks substantial long-term energy reserves and therefore relies on specialized mechanisms to regulate nutrient delivery to specific cell types. One such mechanism is the direct transfer of metabolites between neurons and glial cells such as astrocytes, oligodendrocytes (OLs), and oligodendrocyte progenitor cells (OPCs). A representative example is the glutamate-glutamine cycle, where glutamate released during synaptic activity is taken up by astrocytes and converted into glutamine before being shuttled back to neurons (*1, 2*). The process is crucial for maintaining neurotransmitter balance and allows neurons to conserve energy by recycling neurotransmitter components (*3*). Astrocytes and OLs also provide neurons with important metabolic substrates, such as alanine (*4*) and lactate (*5*), to sustain axonal energy demands. In turn, neurons support glial metabolism by synthesizing cholesterol and supplying it to OPCs, a process that facilitates the repair of demyelinated lesions (*6*). Another well-characterized example of metabolite transfer in the CNS involves N-acetylaspartate (NAA), which is exported from neurons to OLs. Within OLs, NAA is hydrolyzed to acetate and aspartate, providing essential precursors for lipid biosynthesis (*7*). These types of metabolic partnerships between cell types are integral to brain homeostasis and their disruption is a pathological feature of numerous CNS disorders including multiple sclerosis (*8*), inflammatory pain (*9*), stroke (*10*), and dementia (*11*). An improved understanding of metabolic interactions in the brain may thus reveal unexplored therapeutic targets for the treatment of neurological diseases.

A major barrier to advancing our knowledge of metabolic partnerships between cell types within organs is the lack of tools that can track nutrient exchange directly in complex tissue environments. Currently, most of the evidence supporting metabolite transfer between neighboring cells is indirect, lacking empirical measurements of metabolite secretion and uptake within the CNS. For instance, single-cell RNA sequencing (scRNA-seq) has been employed to infer biochemical interactions (*12*), but gene expression profiles do not necessarily reflect protein activities. A more direct strategy to assess nutrient secretion and uptake is to apply metabolomics. The complication is that standard metabolomics protocols are designed for bulk tissue, where signals from specific cell types cannot be deconvolved. Although spatial metabolomics can be applied to image tissues, even with resolutions as fine as 5-10 microns, it is challenging to determine what metabolites originate from which cells (*13*). Moreover, regardless of whether metabolites can be localized to individual cells or not, their provenance remains unresolved. That is, the presence of a metabolite within a cell does not necessarily indicate that it was synthesized there, as it may have originated from a neighboring cell and been transferred prior to the analysis.

To overcome these obstacles that have limited our understanding of metabolic collaborations between neighboring cells, we introduce a technology that we refer to as cell-specific labeling (CSL). In brief, we engineered transgenic mice to express cellobiose-metabolizing enzymes specifically in neurons. Cellobiose is a disaccharide made up of two glucose units linked by a β (1→4) glycosidic bond. Non-engineered mammalian cells are unable to utilize cellobiose. When transgenic mice are administered ^13^C-labeled cellobiose, ^13^C isotopes are exclusively loaded into neurons. The export of neuron-derived metabolites to other cell types results in the appearance of ^13^C isotopes in non-neuronal pathways, providing an experimental readout of nutrient transfer. Using liquid chromatography/mass spectrometry (LC/MS) to perform untargeted metabolomics, we mapped the fate of ^13^C-labeled cellobiose in the brain tissue of transgenic mice across multiple conditions. Our analysis led to the discovery that *myo*-inositol is transferred from neurons to OPCs to support myelin repair following injury.

## Results

### Expression of cellobiose-utilizing genes in neurons to track cell-specific metabolism

To study metabolite transfer in the brain, we used the piggyBac transposon system (*14*) to express two fungal genes from *Neurospora crassa*: the cellobiose/cellodextrin transporter *cdt-1* and the β-glucosidase *gh1-1*. The former allows efficient uptake of cellobiose and the latter cleaves its β (1→4) glycosidic bond to produce two glucose molecules (**fig S1A**). The expression of both transgenes is driven by the neuron-specific synapsin 1 (*Syn1*) promoter (*15*), resulting in the simultaneous expression of CDT-1 and GH1-1 in neurons. Our strategy is to administer ^13^C_12_-cellobiose to transgenic mice and then track the fate of its isotopic labels in bulk brain tissue at the comprehensive level by using untargeted metabolomics (*16*). To identify potential metabolite candidates that could be transferred between cell types, the fate of the isotopic labels must be compared between two conditions. In the first condition, neurons are biochemically coupled to glial cells. Here, isotopically labeled metabolites can be present in multiple cell types. Labeled metabolites are present in neurons because neurons utilize ^13^C_12_-cellobiose directly. Labeled metabolites can also be present in glial cells because cellobiose-derived products are transferred from neurons and further metabolized. Data from the first condition (transfer-ON) are then compared to data from a second condition, where the biochemical coupling between neurons and glial cells is disrupted by a perturbation such as a disease state or drug treatment (transfer-OFF). Specifically, we look for metabolites whose isotopic labeling decreases in the second condition relative to the first as this either reflects: (i) the loss of a glial cell’s ability to take up a cellobiose-derived product and process it further, or (ii) reduced neuronal production of a cellobiose-derived metabolite due to the loss of communication with glia. Metabolites with decreased isotopic labeling in the transfer-OFF condition relative to the transfer-ON condition represent leads that could be transferred from neurons to glia (**Fig. 1A**). They must be subjected to further testing for validation As a first step to establishing our system, we confirmed *cdt-1* and *gh1-1* expression in genetically engineered mice. We refer to transgenic animals positive for both *cdt-1* and *gh1-1* transgene expression as Syn^CSL^ mice. Germline transmission from founder Syn^CSL^ mice was confirmed with positive PCR results for both *gh1-1* and *cdt-1*, while non-transgenic littermate mice (Syn^WT^) were negative for both genes (**Fig. 1B**). Purified primary cortical neurons and whole brain from Syn^CSL^ mice expressed mRNA for both transgenes, whereas wild-type C57BL/6 mice (designated as WT) lacked expression of both *cdt-1* and *gh1-1* (**Fig. 1C**). Transgenic mice are healthy, show similar growth rates to WT mice, and have normal brain architecture as confirmed with staining for myelin, neurofilament-H, and glial fibrillary acidic protein (GFAP) (**Fig. 1, D and E**).

**Fig. 1.**
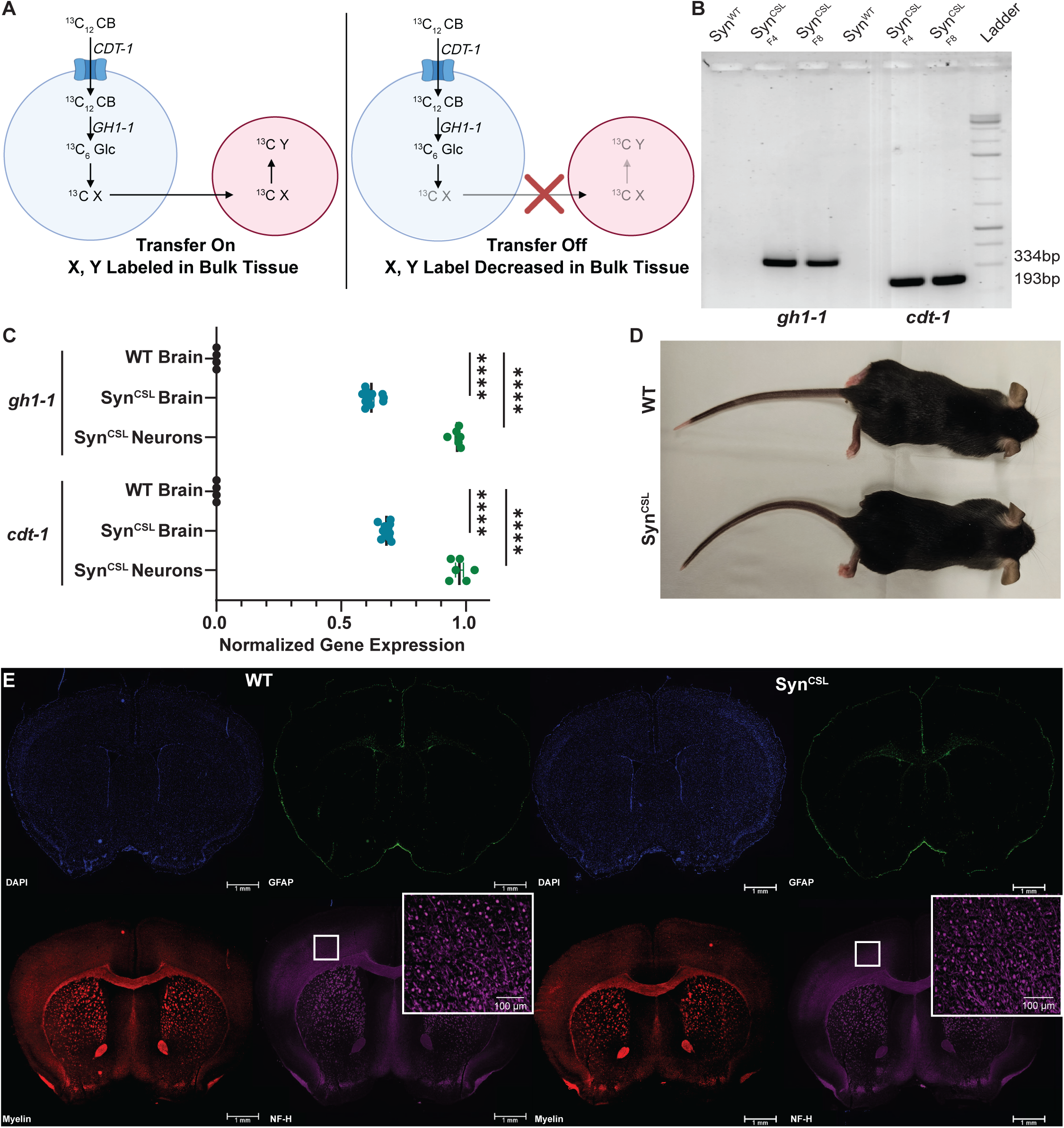
Cell-specific labeling to track metabolite transfer in vivo. **(A)** Experimental design for cell-specific labeling experiments. Expression of *cdt-1* and *gh1-1* permits the uptake and processing of cellobiose (CB) to glucose (Glc). The interruption of transfer through pathology results in differential labeling in bulk tissues. Image made in Biorender. **(B)** PCR results for *cdt-1* and *gh1-1* in genomic DNA from Syn^CSL^ mice or littermate controls (Syn^WT^) at generations 4 and 8. **(C)** Real-time quantitative RT-PCR of transgene expression in brain tissue and primary neurons. Expression normalized with maximum at 1 and minimum at 0. Data represented as mean±SEM. Significance determined via one-sample t-test (control brain, n=4; Syn^CSL^ brain, n=11; Syn^CSL^ neurons=6 animals). **(D)** Representative image of a Syn^CSL^ mouse compared to a C57BL/6J control mouse (WT). **(E)** Immunofluorescence of brains from WT and Syn^CSL^ mice stained for nuclei (DAPI-blue), astrocytes (GFAP-green), myelin (MBP-red), and neurons (neurofilament-H-purple). Inset shows magnified region of neurofila-ment-H staining. Scale bars=1 mm, main image; 100 µm, inset. (* p<.05, ** p<.01, ***p<.001, ****p<.0001).

To demonstrate the functionality of the CDT-1 transporter, we monitored the kinetics of cellobiose uptake by LC/MS in primary cortical neurons. We first confirmed the purity of the primary cultures by immunohistochemical staining with two neuronal markers (anti-NeuN and anti-βIII tubulin) and GFAP to mark astrocytes. The majority of our cells were neurons, with few contaminating astrocytes detected (**fig. S1B**). We then incubated Syn^CSL^ neurons with either 25 mM glucose or 12.5 mM cellobiose and measured the uptake of each substrate from the media over 11 days in culture. The cleavage of cellobiose by GH1-1 produces two glucose molecules. Accordingly, glucose-free media supplemented with 12.5 mM cellobiose contains the same amount of glucose equivalents as media supplemented with 25 mM glucose. In line with the critical role of glucose metabolism in neuronal function (*17*), we observed robust glucose uptake, with ∼30% of the glucose in media depleted after 11 days (**Fig. 2A**). When neurons were cultured in glucose-free media containing cellobiose, cells consumed cellobiose at a rate similar to that of glucose, but depleted only ∼12% of cellobiose from the media (**Fig. 2A**). Although the fractions of glucose and cellobiose depleted were different, the values are consistent with a comparable demand for glucose equivalents. Next, we tested the ability of primary neurons to survive on cellobiose as the sole carbon source by culturing cells for 7 days in 0.5 mM glutamine alone, 0.5 mM glutamine with the addition of 25 mM glucose, or 0.5 mM glutamine with the addition of 12.5 mM cellobiose. Primary Syn^CSL^ neurons cultured in media without a carbohydrate source (glutamine alone) were unable to survive and displayed a 90% reduction in viability (**Fig. 2B**). We also observed a similar decrease in viability when non-transgenic Syn^WT^ neurons, lacking expression of CDT-1 and GH1-1, were cultured with 12.5 mM cellobiose as the sole carbohydrate source (**fig. S1C**). However, the viability of Syn^CSL^ neurons cultured with glutamine and either 25 mM glucose or 12.5 mM cellobiose was not significantly different, indicating that transgenic neurons are able to use either of these two carbohydrates to support cell survival (**Fig. 2B**). While Syn^CSL^ neurons maintained their viability when cultured with cellobiose, they did exhibit a reduction in their basal oxygen consumption rate (OCR) when using cellobiose as substrate compared to glucose. In contrast, mitochondrial respiration was absent in Syn^WT^ cells given cellobiose as their only carbohydrate source (**Fig. 2C**). These results indicate that CDT-1 and GH1-1 are functionally expressed in Syn^CSL^ neurons, allowing them to take up cellobiose and utilize it as a carbon source to support mitochondrial respiration and survival.

**Fig. 2.**
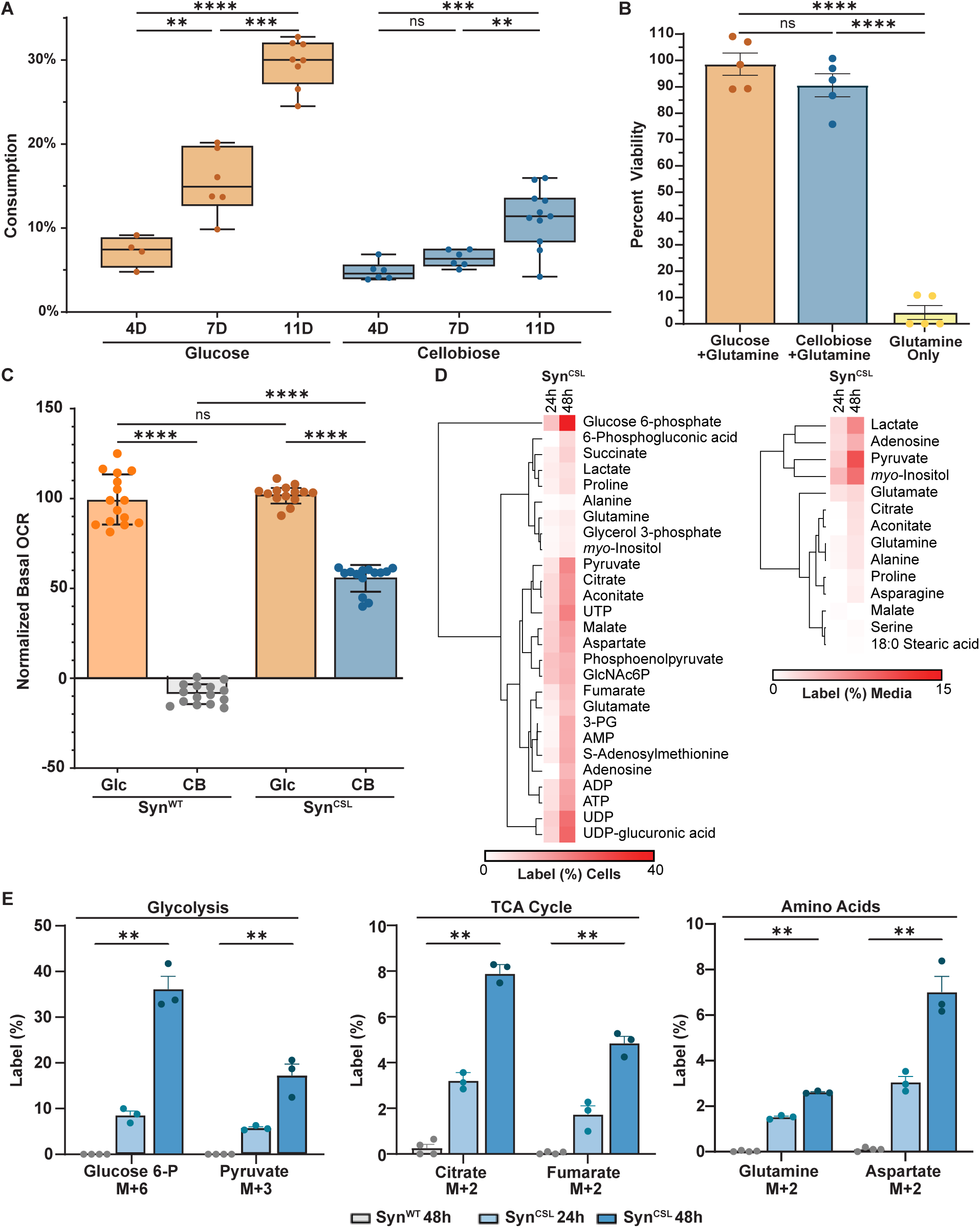
Syn^CSL^ neurons utilize cellobiose as a carbon source *in vitro*. **(A)** Syn^CSL^ neurons uptake glucose and cellobiose *in vitro*. Consumption determined as percent change in media concentration relative to no-cell controls at 4, 7, and 11 days. LC/MS of media from 50k cells/well, error bars=10-90 percentile (glucose 4D, n=4; glucose 7D, cellobiose 4D, and cellobiose 7D, n=6; glucose 11D, n=8; cellobiose 11D, n=11). **(B)** AlamarBlue cell viability assay for 50k neuronal cells/well cultured 7 days in media containing 0.5mM glutamine and either 12.5 mM cellobiose or 25 mM glucose (n=5 wells per group). Data normalized to glucose-treated cultures at 100%. **(C)** Seahorse measurements of basal OCR rates in primary neurons averaged to cell count and normalized to Syn^WT^ cultured with glucose at 100%. Before the assay, 200,000 cells/well were cultured for 3 days in standard media (Syn^CSL^, n=14 wells per treatment; Syn^WT^, n=15 wells per treatment). **(D)** Stable isotope tracing in primary neurons and media after administration of 12.5 mM ^13^C_12_ -cellobiose for 24 or 48 hours. Percent labeling determined by LC/MS and corrected for natural abundance. Euclidean clustering of rows based on average percent labeling for each metabolite. **(E)** Percent labeling of select metabolites in neurons cultured from Syn^WT^ or Syn^CSL^ animals after administration of 12.5 mM ^13^C_12_ -cellobiose for 24 hours (Syn^WT^) or 24 and 48 hours (Syn^CSL^). For this experiment, 2 million cells/well were evaluated (Syn^WT^, n=4 wells; Syn^CSL^, n=3 wells per group (D, E)). Data represented as mean±SEM. Significance determined via Brown-Forsythe and Welch ANOVA with Games-Howell’s post hoc test (A) or ordinary one-way ANOVA with Tukey’s correction (B, C). Significance determined via Kruskal-Wallis ANOVA with Dunn’s test (E) (* p<.05, ** p<.01), *** p<.001, **** p<.0001).

### Cellobiose is a substrate for central carbon metabolism in transgenic neurons *in vitro*

Having demonstrated cellobiose uptake and utilization by Syn^CSL^ neurons, we subsequently aimed to show that expression of CDT-1 and GH1-1 allows isotopic labels from ^13^C_12_-cellobiose to be traced through central carbon metabolism when culturing cells *in vitro*.

Syn^CSL^ or Syn^WT^ neurons were isolated from animals and incubated with 12.5 mM ^13^C_12_-cellobiose for either 24 or 48 hours. Metabolites were then extracted, and samples were analyzed by LC/MS. Substantial labeling was measured in glycolytic metabolites, TCA cycle intermediates, amino acids, and nucleotides from Syn^CSL^ neurons (**Fig. 2D**). These data indicate that the neurons from transgenic mice are able to produce labeled glucose from cellobiose and feed this glucose into both anabolic and catabolic pathways. Similar to intracellular metabolites, we also observed labeling in a smaller number of secreted metabolites, including lactate, pyruvate, and *myo*-inositol (**Fig. 2D**). Following the addition of ^13^C_12_-cellobiose, intracellular glucose 6-phosphate (glucose 6-P) showed approximately 10% labeling after 24 hours and 35% labeling after 48 hours. As expected, lower labeling percentages were observed in downstream metabolites such as M+3 pyruvate, M+2 citrate, M+ 2 fumarate, M+2 glutamine, and M+ 2 aspartate (**Fig. 2E**). As a control, Syn^WT^ neurons were incubated with 12.5 mM ^13^C_12_-cellobiose for 48 hours. No labeling was observed in any metabolites examined from these cells, including glucose 6-P, pyruvate, citrate, fumarate, glutamine, and aspartate (**Fig. 2E**). We also compared the efficiency of labeling after administration of either ^13^C_12_-cellobiose or ^13^C_6_-glucose to cultured Syn^CSL^ neurons over the same time period. Consistent with our basal OCR measurements, we observed lower labeling percentages in metabolites when cells were incubated with ^13^C_12_-cellobiose compared to ^13^C_6_-glucose (**Fig. 2E** and **fig. S1D, E**). Importantly, however, the same sets of metabolites were labeled after incubation with either tracer, revealing that glucose and cellobiose share the same metabolic fate.

### Cell-specific labeling tracks global neuronal contributions to healthy brain metabolism

Having shown that our system works as expected *in vitro*, we next aimed to validate it in *in vivo*. To deliver tracer directly to the brain, we implanted a cannula into the right lateral ventricle of Syn^CSL^ mice and administered either 400 mM ^13^C_12_-cellobiose or 800 mM ^13^C_6_-glucose at a rate of 8 µL/hour with an osmotic pump for 1, 3, 6, 12, 18, or 24 hours (**Fig. 3A**). Our rationale for delivering isotopic tracers directly into the CNS is two-fold. First, the continuous infusion of isotope allows us to achieve isotopic steady-state in pathways such as glycolysis (*18*). Second, the direct delivery of cellobiose to the brain prevents its potential degradation by the microbiome, which could complicate our interpretation of labeling results (*19*). Imaging cellobiose-infused brains by mass spectrometry showed the predicted pattern of substrate diffusion, with higher concentrations of cellobiose observed more proximal to the lateral ventricle where the cannula was placed (**fig. S2A**). Our strategy to continuously deliver stable isotope through the intracerebroventricular (ICV) route did not perturb metabolic homeostasis, as the infusion of cellobiose or glucose did not significantly change the levels of metabolites in the brain (**fig. S2B**). Additionally, as expected, no labeling was detected in WT mice administered ^13^C_12_-cellobiose (**fig. S2C**). These data demonstrate that CDT-1 and GH1-1 expression are required for the brain to utilize cellobiose as a nutrient *in vivo*. We then examined the labeling kinetics of glycolytic intermediates in brain tissue following the infusion of ^13^C_12_-cellobiose or ^13^C_6_-glucose. Over the course of 24 h when the tracers were provided, the labeling of glycolytic intermediates was not significantly different between ^13^C_12_-cellobiose and ^13^C_6_-glucose experiments. We also note that pseudo isotopic steady state occurred by 24 h with both tracers (**Fig. 3B**). In contrast to glycolysis, downstream metabolic pathways such as the TCA cycle showed a significant reduction in labeling from ^13^C_12_-cellobiose compared to ^13^C_6_-glucose (**Fig. 3C, D and fig. S2D**). These results indicate that glycolytic labeling from bulk brain tissue is primarily driven by neurons. In other words, the extent to which glucose fuels glycolysis in non-neuronal cell types throughout the brain is relatively small. The disparity in the fraction of TCA cycle metabolites that are labeled when using ^13^C_12_-cellobiose and ^13^C_6_-glucose tracers, on the other hand, suggests that neurons oxidize a considerable amount of metabolic substrates other than glucose. This result is in line with previous studies, supporting the idea that mitochondrial metabolism in neurons is supported by nutrients transferred from glial cells (*4, 20, 21*).

**Fig. 3.**
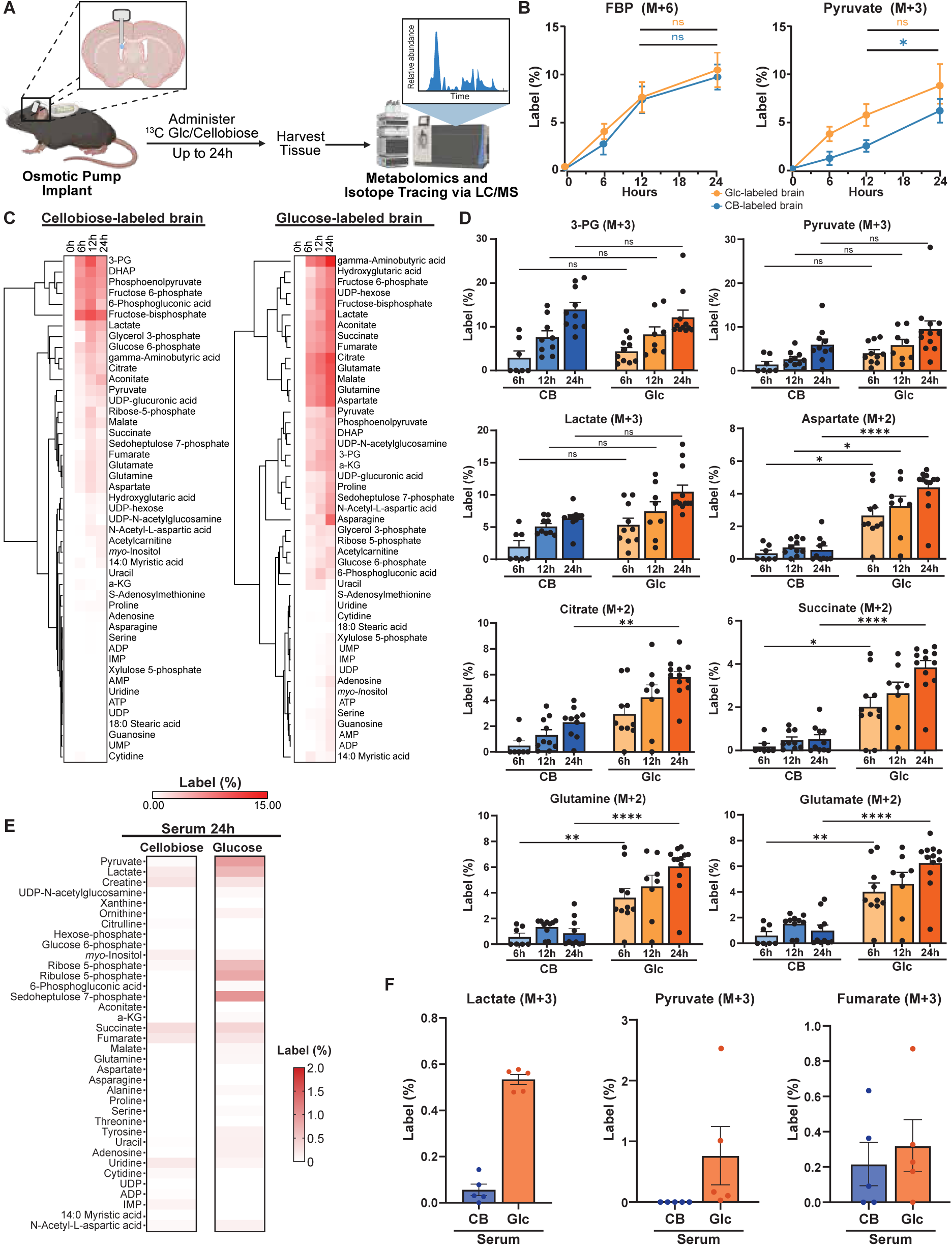
Cellobiose is utilized as fuel source in vivo by Syn^CSL^ mice. **(A)** Experimental scheme of osmotic pump implantation for isotope tracing. Image made in Biorender. **(B)** Time course of percent labeling of M+6 FBP and M+3 pyruvate in brains during ICV administration of ^13^C_6_ -glucose or ^13^C_12_-cellobiose. Mice were treated with 800 mM ^13^C_6_ -glucose or 400 mM ^13^C_12_-cellobiose at 8 µl/hour for 0, 6, 12, or 24 hours. Symbols represent mean±SEM. Significance of 12 vs. 24 hours determined via Mann-Whitney test with Holm-Sidak correction. Significance of glucose vs cellobiose treatment determined via functional data analysis (p-value FBP=.997; p-value pyruvate=.279). **(C)** Heatmap of total average percent labeling in metabolites from brains after administration of 800 mM ^13^C_6_ -glucose or 400 mM ^13^C_12_ -cellobiose. Euclidean clustering of rows is based on average. **(D)** Percent labeling over time for M+3 3-PG, M+3 pyruvate, M+3 lactate, M+2 aspartate, M+2 citrate, M+2 succinate, M+2 glutamate, and M+2 glutamine in whole-brain sections. Significance determined via Kruskal-Wallis test with Dunn’s correction (CB 0h, n=6 mice; Glc 0h, n=15; CB 6h and Glc 12h, n=8; Glc 6h and CB 24h, n=10; CB 12h, n=9; Glc 24h, n=12 animals). (B, C, D)). **(E)** Heatmap of total percent labeling of metabolites in the serum of mice administered ^13^C -glucose or ^13^ -cellobiose for 24 hours. **(F)** Percent labeling of M+3 lactate, M+3 pyruvate, and M+3 fumarate in serum. (n=5 mice per condition, (E, F)). Bars represent mean±SEM (D, F). Percent labeling determined by LC/MS and corrected for natural abundance (B, C, D, E, F). (* p<.05, ** p<.01), *** p<.001, **** p<.0001).

An intriguing application of our cell-specific labeling technology is that we can examine the metabolic contribution that neurons make to systemic circulation for the first time. When ^13^C_12_-cellobiose is infused directly into the ventricles of the brain, it is exclusively loaded into neurons. Other ^13^C-labeled metabolites can only be produced by neuronal metabolism, either directly by biochemical pathways in neurons or indirectly through the modification of neuron-derived metabolites in other cell types. We provided ^13^C_12_-cellobiose or ^13^C_6_-glucose via ICV infusion for 24 hours and analyzed labeled metabolites that appeared in serum. TCA cycle intermediates, nucleotides, and amino acids all displayed labeling in serum from both tracers (**Fig. 3E, F**). As an example, although labeling was observed in serum lactate following administration of both ^13^C_12_-cellobiose and ^13^C_6_-glucose, labeling in pyruvate was only measured in serum after infusion of ^13^C_6_-glucose. These data indicate that a fraction of circulating lactate and pyruvate originate from glucose in the brain. The results suggest that most of the brain-derived pyruvate in the circulation originates from glial cells, whereas at least ∼10% of brain-derived lactate tracks back to neurons. Our findings highlight the application of Syn^CSL^ mice to gain a richer understanding of the cellular provenance of circulating metabolites, which could potentially be leveraged to develop blood biomarkers of brain pathology.

### Cell-specific labeling identifies *myo*-inositol transfer from neurons to OPCs

We aimed to use cell-specific labeling to identify previously unknown nutrient transfers between neurons and glia during CNS pathology. To that end, we utilized cuprizone (biscyclohexanone oxaldihydrazone, CPZ) to induce demyelination as a prototypic model of multiple sclerosis (*22, 23*). CPZ is a copper chelator and mitochondrial toxin that selectively induces cell death in mature, myelinating OLs, leading to myelin loss in white matter regions such as the corpus callosum. This model closely resembles the OL pathology seen in primary demyelinating disorders and is frequently used to study therapies aimed at promoting remyelination (*24–28*). CPZ exhibits strain- and age-specific pathology, with C57BL/6 mice being the most widely used (*29*). Although we backcrossed our transgenic mice to the C57BL/6 background for at least seven generations, we wanted to confirm that the severity and kinetics of demyelination in Syn^CSL^ mice were similar to age-matched C57BL/6 mice. Eight-week-old mice were fed 0.25% CPZ (w/w) in their diet for 6 weeks. In a second set of mice, CPZ was withdrawn after six weeks of feeding and animals were allowed to undergo remyelination for 2 weeks or 4 weeks (**Fig. 4A**). Immunohistochemical analysis of demyelination and gliosis showed that myelin loss and astrocyte activation were not different between CPZ-treated Syn^CSL^ mice and controls, which included both Syn^WT^ littermates and C57BL/6 mice. In all cases, peak demyelination was observed after 6 weeks of feeding (**Fig. 4B, fig. S3A**). Coinciding with the loss of myelin, mice displayed robust recruitment of OPCs to demyelinating lesions (**fig. S3B**). Following withdrawal of CPZ from the diet, Syn^CSL^ mice also had similar kinetics of remyelination (**Fig. 4B**) compared to controls (*30*).

**Fig. 4.**
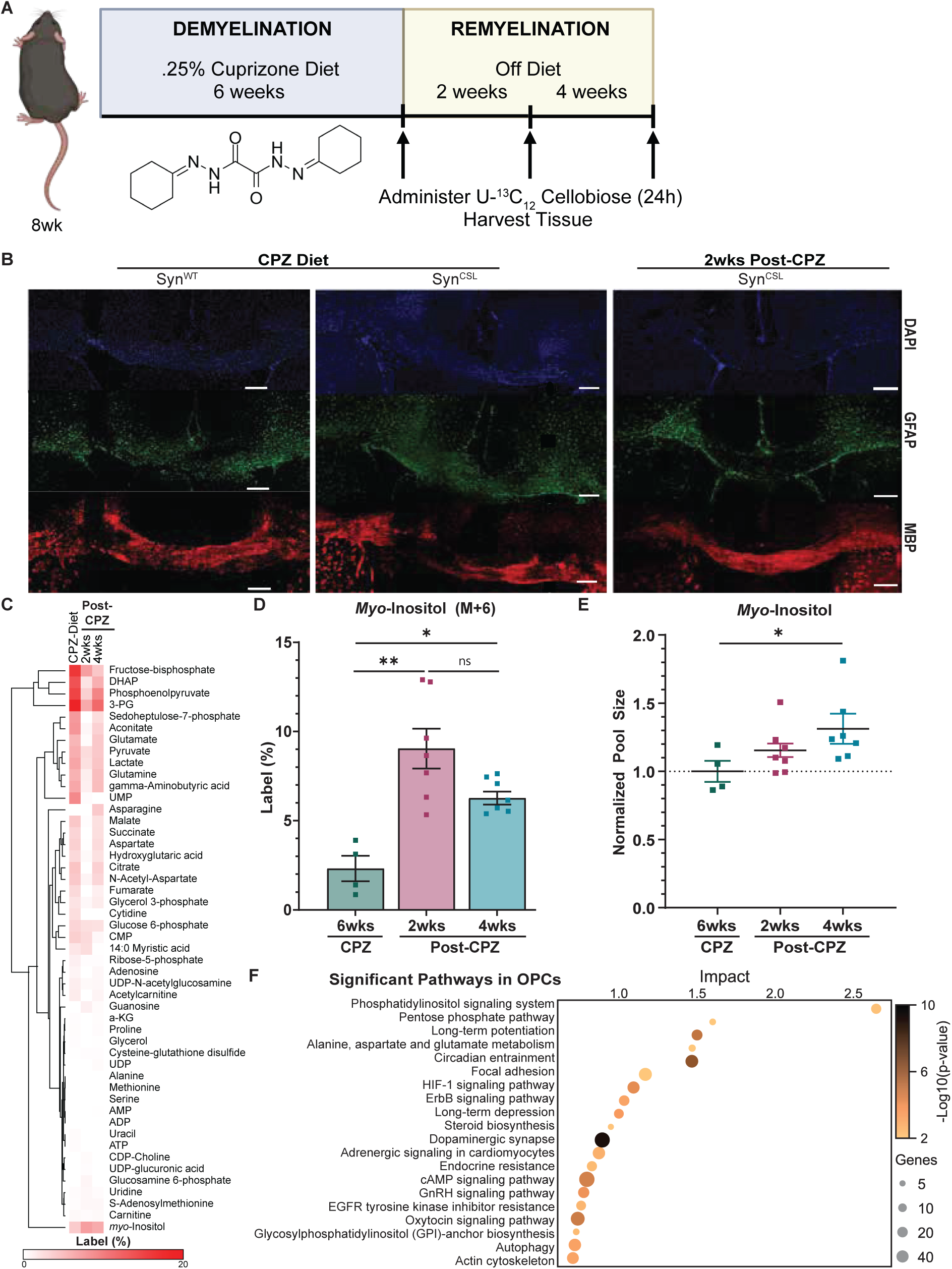
Remyelination stimulates *myo*-inositol transfer between neurons and OPCs. **(A)** Scheme of CPZ intoxication and time line for isotope tracing. Image made in Biorender. **(B)** Representative immunofluorescence images of brains from Syn^WT^ and Syn^CSL^ mice at week 6 of CPZ treatment or 2-weeks post-CPZ diet stained for nuclei (DAPI-blue), astrocytes (GFAP-green), and myelin (MBP-red). Scale bars are 250 µm. **(C)** Heatmap of total average percent labeling from brains of Syn^CSL^ mice after 6 weeks of CPZ diet and 2- and 4-weeks post-CPZ diet. Euclidean clustering of rows based on average percent labeling for each metabolite. **(D)** Percent labeling of M+6 *myo*-inositol in the corpus callosum of mice treated with ^13^C_12_-cellobiose for 24 hours alongside either 6 weeks of CPZ diet or 2 or 4 weeks post-CPZ diet. Bars represent mean, error bars represent SEM. Percent labeling determined by LC/MS and corrected for natural abundance. **(E)** Scatterplot of *myo*-inositol pool size in the corpus callosum normalized to 6 weeks CPZ diet. Lines represent mean±SEM. (CPZ diet, n=4 animals; 2wks and 4wks post-CPZ, n=7 animals. (C, D, E)). **(F)** Dot plot of significantly enriched pathways (FDR<0.05) from snRNAseq in OPCs during remyelination vs demyelination via CPZ intoxication. Dot size represents the number of genes represented, color represents -log10(p-value). Significance determined via Brown-Forsythe and Welch ANOVA with Games-Howell’s post-hoc (D) or Dunnett’s correction (E) (* p<.05, ** p<.01), *** p<.001).

After establishing that disease pathology was conserved in transgenic animals, we gave Syn^CSL^ mice an ICV infusion of ^13^C_12_-cellobiose for 24 hours. In one experiment, cellobiose tracer was administered after 6 weeks of CPZ feeding, when demyelination was prominent. In separate experiments, ^13^C_12_-cellobiose was administered following a 2- or 4-week withdrawal period over which time CPZ was removed from the food to stimulate remyelination. Isotopic labeling was measured in the corpus callosum of all samples by using LC/MS. We reasoned that remyelination would result in increased nutrient transfer between neurons and OPCs to support myelin production, representing the transfer-ON state. Conversely, we hypothesized that demyelination would disrupt transfer between neurons and OPCs, representing the transfer-OFF state (**Fig. 1A**). Within the corpus callosum, we observed alterations in the labeling of glycolytic metabolites, TCA cycle intermediates, neurotransmitters, and NAA over the time course of demyelination and remyelination. The majority of the metabolites found to incorporate ^13^C showed their highest level of labeling during demyelination. Labeling initially decreased after withdrawal of CPZ for 2 weeks, but increased again by 4 weeks of CPZ withdrawal (**Fig. 4C**). In contrast to this trend, changes in *myo*-inositol labeling were unique (**Fig. 4C, D**). In addition to having elevated concentrations, *myo*-inositol labeling was higher at both the 2 week and 4 week remyelinating time points relative to the demyelinating time point (**Fig. 4D, E**). An increase in the fraction of labeled *myo*-inositol during remyelination suggested that it is part of an intercellular transfer process (**Fig. 1A**).

To gain insight into how *myo*-inositol metabolism is partitioned between neurons and OPCs, we complemented our cellobiose tracing results with single-nucleus RNA sequencing (snRNA-seq) data from a previously published study examining CPZ intoxication, which provided cell-type specific changes in transcription during demyelination and remyelination (*31*). The snRNA-seq data showed that pathways downstream of *myo*-inositol were enriched in OPCs, but not neurons during remyelination (**fig. S3C, D**). Indeed, when comparing snRNA-seq data from the demyelination and remyelination states, the top enriched metabolic pathway (FDR < 0.05) in OPCs was the phosphatidylinositol (PI) signaling system (**Fig. 4F**). These findings support a mechanism in which *myo*-inositol production is increased in neurons after CPZ withdrawal to support transfer to OPCs, where *myo*-inositol is used to synthesize PI lipids for myelin repair.

To gain more evidence for this model, we performed four additional analyses. First, we measured the expression of genes involved in *myo*-inositol production and utilization within the corpus callosum during demyelination and remyelination. Two genes associated with the production of *myo*-inositol from glucose, inositol monophosphatase 1 (*Impa)* and inositol-3-phosphate synthase (*Isyna)*, were decreased during CPZ-induced demyelination but increased in response to remyelination. A similar expression pattern was measured for CDP-diacylglycerol-inositol 3-phosphatidyltransferase (*Cdipt*) and phosphatidylinositol glycan anchor biosynthesis class A (*Piga*) (**Fig. 5A** and f**ig. S3E**). The former encodes an enzyme in PI lipid biosynthesis, while the latter is required for PI-dependent membrane functions. The data are consistent with increased production of *myo*-inositol in the corpus callosum during remyelination. Second, to confirm that *myo*-inositol derived from ^13^C_12_-cellobiose is incorporated into the polar head group of PI lipids, we evaluated corpus callosum tissue after CPZ withdrawal by LC/MS. Specifically, using corpus callosum tissue from animals administered ^13^C_12_-cellobiose, we fragmented PI lipids and verified the presence of six ^13^C labels in the head group (**fig. S3F**). Third, we demonstrated that neurons synthesize *myo*-inositol *in vitro* and secrete it into the culture medium (**fig. S3G-I**). And, fourth, we superimposed metabolites labeled from ^13^C_12_-cellobiose during CPZ withdrawal on the same pathway map as genes having increased expression during remyelination. Using snRNA-seq data, separate pathway maps were constructed for genes uniquely increased in neurons and genes uniquely increased in OPCs. When the tracer and snRNA-seq results converge in one map but not the other, it suggests that the labeled metabolite is transformed in that specific cell type. The analysis indicates that neuron-derived *myo*-inositol is incorporated into PI lipids within OPCs, rather than PI lipids being synthesized within neurons and transferred to OL precursor cells (**Fig. S3C,D**).

**Fig. 5.**
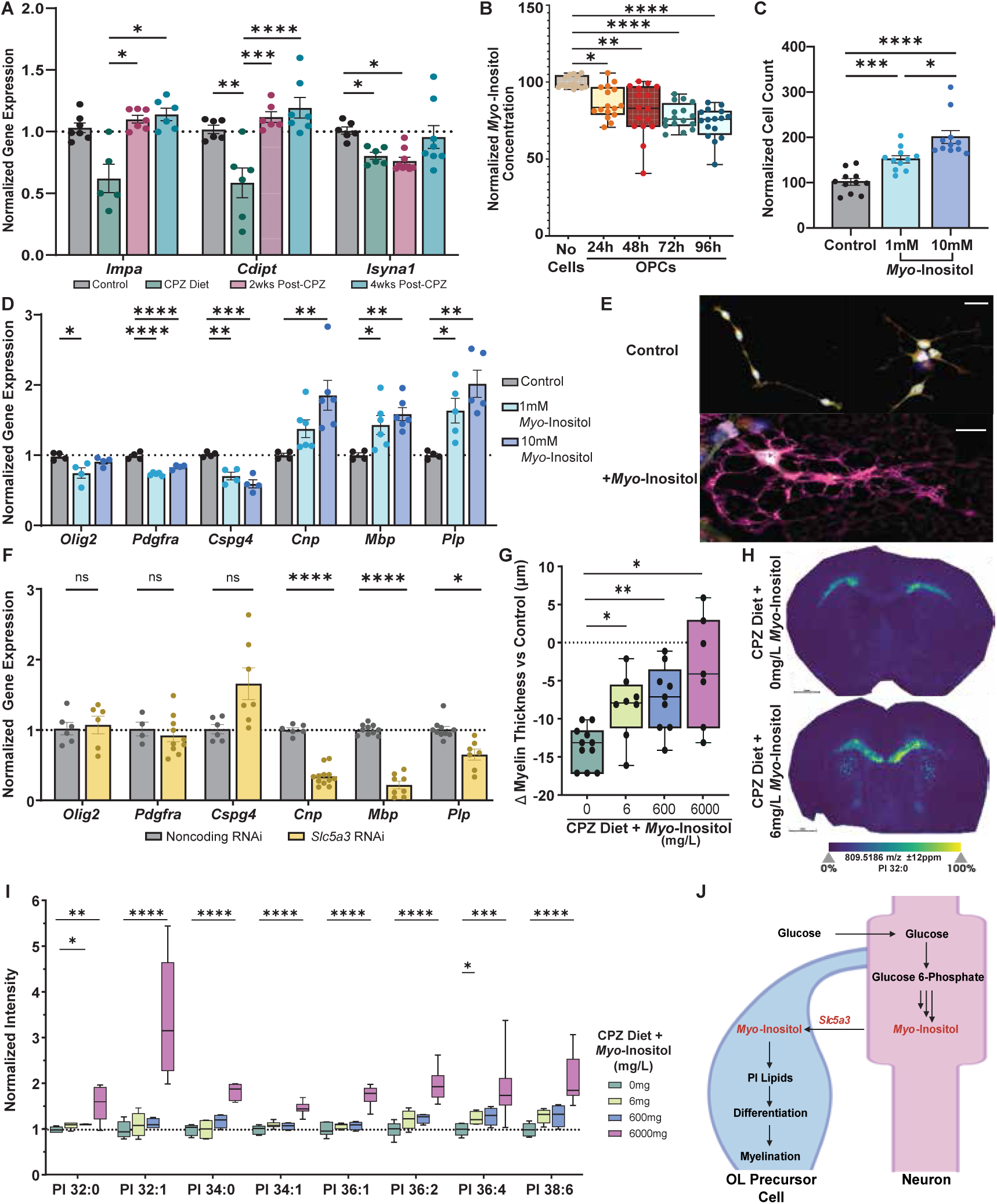
***Myo*-inositol increases OPC differentiation through SLC5A3 and promotes myelin repair. (A)** Normalized gene expression of inositol-associated metabolic genes in the corpus callosum of mice administered 6 weeks of standard diet, 6 weeks of CPZ diet, or 6 weeks of CPZ diet followed by 2 or 4 weeks on standard diet. Data normalized to expression of control mice fed standard diet by using the *Gapdh* housekeeping gene (control, n=6; CPZ-diet, n≥5 mice; 2wks post-CPZ, n≥6; 4wks post-CPZ, n≥6). **(B)** Box-and-whiskers plot of *myo*-inositol concentration in OPC media after the addition of 10 mM *myo*-inositol. Data normalized to no-cell controls. Per well, 50,000 OPCs were plated. (no-cell control, n=14 wells; 24, 48, 72, and 96H, n=16 wells). **(C)** Cell proliferation of primary OPCs (200,000 cell/well) after culture in 0, 1, or 10 mM *myo*-inositol media for seven days. Data normalized to 0 mM control (n=11 wells per group). **(D)** Normalized gene expression of OPCs (200,000 cell/well) after culture in 0, 1 mM, or 10 mM *myo*-inositol containing media. Data normalized to controls cultured in 0 mM *myo*-inositol by using the *Gapdh* housekeeping gene (n≥4 wells per group). **(E)** Representative immunofluorescence image of OPCs cultured for 7 days in media containing 0 mM or 1 mM *myo*-inositol and stained for nuclei (DAPI-blue), OPC markers (CD140-red, vimentin-green), and oligodendrocyte marker (MBP-pink). Scale bars represent 25 µm. **(F)** Normalized gene expression of OPCs (200,000 cells/well) cultured in media containing 5 mM *myo*-inositol and treated with *Slc5a3* or noncoding RNAi. Data normalized to noncoding RNAi control by using the *Gapdh* housekeeping gene (n≥4 wells per condition). Significance determined via multiple T-tests with Holm-Sidak correction. **(G)** Box-and-whiskers plot of the change in myelin thickness of CPZ-fed mice with or without *myo*-inositol supplementation relative to control mice fed standard diet. Myelin thickness measured by immunofluorescence for MBP (0 mg/L, n=11; 6 mg/L, n=8; 600 mg/L, n=9; 6000 mg/L, n=7 animals). **(H)** MALDI images of PI 16:0/16:0 [M-H]^-^ adduct in mice fed a CPZ diet without (top) or with (bottom) 6 mg/L *myo-*inositol supplementation. Images are normalized on the same intensity scale, scale bars represent 1 mm. **(I)** Normalized intensity of pool sizes of PIs as measured by LC/MS in the corpus callosum of mice fed a CPZ diet for 6 weeks with or without *myo*-inositol supplementation (0 mg and 6 mg/L, n=7; 600 mg/L, n=4; 6000 mg/L, n=10 animals). Data normalized to 0 mg/L. **(J)** *Myo*-inositol production by neurons and transfer to OPCs supports PI lipid synthesis and myelination. Bars represent mean±SEM (panels A, C, D, F). Error bars represent min to max (B, G, I). Significance determined via one-way ANOVA with Tukey’s correction (A, C). Significance determined with Brown-Forsythe and Welch ANOVA with Dunnett’s correction (B, D, G, I). (* p<.05, ** p<.01), *** p<.001, **** p<.0001).

### *Myo*-inositol transfer from neurons to OPCs supports remyelination

We speculated that *myo*-inositol export from neurons may support OPC differentiation and myelin production. To explore the functional role of *myo*-inositol in OPCs, we first confirmed that primary murine OPC cultures take up *myo*-inositol from the culture medium. When incubated with 10 mM *myo*-inositol, a concentration that is consistent with the 2-15 mM levels measured from the brain (*32*), we observed significant uptake over the course of 96 hours (**Fig. 5B**). Cellular uptake of *myo*-inositol by OPCs was associated with significantly increased cell proliferation, and this effect was dose dependent (**Fig. 5C**). A key step in myelin repair is the migration of OPCs to sites of damage, followed by OPC proliferation and differentiation into mature myelinating OLs (*33–35*). We postulated that in addition to supporting proliferation, *myo*-inositol might also stimulate the differentiation of OPCs. Indeed, the expression of genes associated with OPCs and pre-myelinating OLs (*Cspg4*, *Pdgfra*, and *Olig2*) were decreased after *myo*-inositol treatment, while genes involved in myelin synthesis and expressed by mature, myelinating OLs (*Cnp*, *Mbp*, and *Plp*) were increased in response to treatment (**Fig. 5D**). Similar changes in proliferation and myelin-associated gene expression were also observed when OPCs were incubated with conditioned media from primary neuronal cultures (**fig. S4A, B**). Consistent with our gene expression analysis, OPCs treated with *myo*-inositol displayed a morphology resembling mature, myelinating OLs with increased myelin basic protein (MBP) staining (**Fig. 5E**).

The uptake of *myo*-inositol into cells is mediated by three members of the SLC family of transporters. SLC5A3 (SMIT1) and SLC5A11 (SMIT2) are characterized as Na^+^/*myo*-inositol cotransporters (*36, 37*). A third transporter, HMIT (SLC2A13) transports *myo*-inositol along with a proton (*38*). The function of these three transporters in the brain have been primarily investigated in neurons and astrocytes (*37, 39*), but some evidence suggests myelin injury can alter the expression of SLC5A3 in OPCs (*40*). Thus, we sought to test whether knockdown of *Slc5a3* in primary OPCs would impact *myo*-inositol-induced differentiation. We first confirmed that *Slc5a3* is expressed in purified murine OPCs and that we could effectively knock it down with siRNA (**fig. S4C**). We then verified that knockdown of *Slc5a3* prevented *myo*-inositol uptake by OPCs (**fig. S4D**). Next, we determined the impact of *Slc5a3* knockdown on myelin-associated gene expression. In line with our earlier results where *myo*-inositol supplementation stimulated expression of genes involved in myelin synthesis (*Cnp*, *Mbp*, and *Plp*), knockdown of *Slc5a3* significantly reduced the expression of these same genes in primary OPCs compared to controls (**Fig. 5F**). Here, controls were OPCs treated with noncoding RNAi and *myo*-inositol. The results point to an unexplored role for SLC5A3 in *myo*-inositol uptake by OPCs to support myelin production and cellular differentiation.

Finally, we wanted to test whether dietary supplementation with *myo*-inositol could rescue deficient nutrient transfer during demyelination. To explore this question, we fed mice 0.25% CPZ (w/w) for six weeks and added *myo*-inositol in the drinking water at doses ranging from 6 mg/L to 6000 mg/L starting after week three of CPZ treatment. Supplementation at these doses led to an increase in circulating *myo*-inositol levels as well as increased *myo*-insitol at the sites of demyelination (**fig. S4E, F**). The thickness of the corpus callosum was subsequently measured at week six in the following groups: (i) CPZ-treated animals, (ii) CPZ-treated animals receiving *myo*-inositol in the drinking water, and (iii) age-matched controls receiving neither CPZ nor *myo*-inositol. As expected in animals fed CPZ, there was a significant reduction in corpus callosum thickness compared to controls due to the CPZ-induced loss of myelinating OLs (*23, 41*). In contrast, animals that received *myo*-inositol in the drinking water for 3 weeks during CPZ feeding were able to maintain myelin in the corpus callosum. With dietary supplementation, the thickness of the corpus callosum was significantly increased compared to CPZ-treated animals and this effect was dose dependent (**Fig. 5G** and **fig. S4G**). Additionally, transcription of *Slc5a3* was modulated by both demyelination and remyelination, with its lowest expression levels in corpus callosum after 6 weeks of CPZ feeding. *Myo*-inositol supplementation for 3 weeks significantly increased *Slc5a3* expression to control levels. During the period of remyelination, after CPZ had been withdrawn for 2 weeks, *Slc5a3* expression was found to be the highest (**fig. S4H**). Based on our other results, we postulated that *myo*-inositol supplementation supported PI lipid metabolism. To investigate this possibility, we analyzed corpus callosum tissue with both LC/MS and mass spectrometry imaging. The data showed that treatment of animals with increasing concentrations of *myo*-inositol (6 mg/L - 6000 mg/L) in the water during CPZ feeding led to increased levels of multiple PI lipids in the corpus callosum (**Fig. 5H, I**). Taken together, our results suggest a model in which neurons synthesize *myo*-inositol from glucose and transfer it to OPCs through the activity of SLC5A3, leading to increased production of PI lipids that support OPC proliferation, differentiation, and myelin production (**Fig. 5J**).

## Discussion

Metabolic coordination between different cell types within an organism is essential to maintaining physiological homeostasis. One way in which coordination is achieved is by transferring metabolites, so that the metabolic output of one cell type is the metabolic input of another. Although this type of biochemical crosstalk is a defining feature of *in vivo* metabolism, our knowledge of which specific substrates are exchanged between various cell types throughout an organism remains largely incomplete due to measurement limitations. To address this critical gap in our understanding of metabolism, here we introduce cell-specific labeling as a metabolomics technology to track biochemical interactions *in vivo*.

Historically, a major challenge of characterizing biochemical crosstalk with metabolomics has been determining the cellular provenance of metabolites. Mass spectrometry provides a readout of what metabolites are present, but it does not directly reveal from which cell type the metabolites originated. As a consequence, to study nutrient transfer by metabolomics, researchers have often evaluated conditioned media from single cell types cultured in isolation (*42*). The idea is to identify metabolites secreted into media by one cell type and subsequently consumed from the media by another. While this workflow has led to important discoveries (*43*), it is fundamentally limited by the non-physiological conditions under which it is performed. The experimental design cannot capture metabolic interactions that require physical proximity, as the different cell types are not in direct contact. Moreover, it does not replicate the bidirectional and dynamic aspects of biochemical crosstalk that occur *in vivo*. Here, we establish cell-specific labeling to track the cellular provenance of metabolites in the intact animal for the first time, thereby enabling nutrient transfer to be studied by metabolomics under physiologically relevant conditions.

As a first demonstration of cell-specific labeling, we engineered transgenic mice to express cellobiose-metabolizing enzymes specifically in neurons. We focus on an application related to demyelination in this work but the mice we established will be broadly useful for studying a wide range of neurodegenerative or neurovascular pathologies in which neuronal metabolism is disrupted. Our genetic system is also adaptable to other cell populations, organ systems, or additional pathologies such as cancer by using cell-type specific promoters to express *cdt-1* and *gh1-1*. We note that constitutive expression of fungal genes in neurons from birth under the control of the *Syn1* promoter did not result in abnormal development, cell toxicity, or inappropriate activation of the immune system. Therefore, it is likely that these genes can be expressed in multiple cell types or target organs without pathology.

Through the application of cell-specific labeling, we discovered that neurons transfer *myo*-inositol to OPCs, stimulating their proliferation and differentiation. *Myo*-inositol is an abundant brain metabolite, acting as both an osmolyte (*44*) and a precursor to PI lipids (*32*). The role of *myo*-inositol in OPCs and myelination, however, has remained largely unexplored. In this study, we demonstrate that *myo*-inositol is derived from glucose in neurons and transferred to OPCs to promote PI lipid production. Although these lipids are a minor component of cell membranes compared to other glycerophospholipids, they have a high turnover rate as their phosphorylation produces important phosphoinositide second messengers including PI(3,4)P_2_, PI(4,5)P_2_, and PI(3,4,5)P_3_ (*45, 46*). The effects of neuron-derived *myo*-inositol in OPCs may therefore be mediated by the production of second messengers, acting as a signal from demyelinated axons to initiate repair. Activation of the phosphoinositide 3-kinase (PI3K)/Akt pathway, which produces PI(3,4,5)P_3,_ has been shown to regulate both OPC survival (*47*) and differentiation (*48*). Additionally, PI(4,5)P_2_ binds to MBP and stabilizes its association with myelin membranes (*49*), a finding consistent with our observation that *myo*-inositol treatment increases MBP protein expression in OPCs. Taken together, our data suggest that *myo*-inositol production in neurons and its transfer to OPCs may serve as a local differentiation signal that promotes myelin formation.

The *myo*-inositol transporter, SLC5A3, plays a critical role in *myo*-inositol uptake by OPCs. While the function of this transporter within the brain has primarily been characterized in neurons (*50*), astrocytes (*51*), and choroid plexus epithelium (*52*), its role in OLs or OPCs remains less well understood. Whole-body deletion of SLC5A3 leads to respiratory failure and death shortly after birth, a phenotype that can be rescued by maternal *myo*-inositol supplementation (*53*). The developmental defects from SLC5A3 deficiency have largely been attributed to altered inositol metabolism in neurons (*37, 53*), but impaired myelination may also contribute to the observed pathologies. Our data show that knockdown of *Slc5a3* in OPCs blocks the pro-myelinating effects induced by *myo*-inositol treatment. In contrast, remyelination following CPZ administration, and to a lesser extent *myo*-inositol supplementation, results in increased expression of *Slc5a3*. Our findings suggest that the recruitment and differentiation of OPCs *in vivo* is associated with upregulation of the SLC5A3 transporter. Further studies are needed to understand the regulation of SLC5A3 expression and activity in OPCs, as well as whether its expression is linked to specific progenitor phenotypes (*54, 55*).

We find that *myo*-inositol supplementation during CPZ feeding increased the efficiency of on-going myelin repair. Previous studies examining the impact of dietary *myo*-inositol supplementation in mood disorders have shown that oral administration of *myo*-inositol increases CNS *myo*-inositol levels in both animal models and humans (*56, 57*). Additionally, though the impact on demyelination was not extensively evaluated, an exploratory study of nine multiple sclerosis patients demonstrated that *myo*-inositol supplementation is well-tolerated at doses up to 1000 mg/day (*58*). In our study, dietary *myo*-inositol supplementation elevated levels of both *myo*-inositol and PI lipid species in the corpus callosum and helped to preserve the integrity of the myelin sheath after CPZ feeding. While CPZ induces myelin injury similar to that observed in multiple sclerosis, it does not replicate the inflammatory microenvironment that is a key characteristic of demyelinating lesions (*28*). That said, current disease-modifying therapies for multiple sclerosis are extremely effective at reducing inflammation but cannot stimulate myelin repair. Thus, manipulating *myo*-inositol levels through dietary supplementation, in combination with existing anti-inflammatory treatments, may offer a promising therapeutic strategy for multiple sclerosis and other demyelinating disorders.

## Materials and Methods

### Animals

Animal work was approved by the Institutional Care and Use Committee at Washington University (protocol 19-0930 and 22-0304). All mice used in this study were at least 8 weeks old at the start of the experiment and experimental groups contained a mix of male and female mice. Mice were housed at a density of 5 animals per cage with *ad libitum* access to standard chow diet (Purina PicoLab Rodent Diet 20), unless otherwise indicated. The colony was maintained in humidity- and temperature-controlled rooms under a 12-hour light-dark cycle and all procedures were performed during the light cycle.

Syn^CSL^ mice were generated by Cyagen Biosciences (Santa Clara, CA, USA) by using piggyBac transposon technology. The coding sequences for the *Neurospora crassa cdt-1* and *gh1-1* genes were optimized for human codon usage. Founder mice were subsequently bred to a C57BL/6 background and genotype confirmed via PCR. Control animals were either littermates that were PCR-confirmed negative for transgene expression or C57BL/6 animals obtained from Charles River Laboratories (Strain # 027).

### Genotyping of Syn^CSL^ Mice

Tissue samples were collected as a 2mm ear punch and DNA was extracted by using the HotSHOT DNA extraction method. In short, 75µl of alkaline lysis buffer (25mM NaOH, .2mM disodium EDTA in water, Sigma Aldrich, St. Louis, MO, USA) was used to submerge tissue. Samples were heated at 95°C for 45 minutes, cooled at 4°C for 15 minutes, and 75µl of neutralizing buffer (40mM Trizma-HCl, Sigma Aldrich, St. Louis, MO, USA) was added and mixed. PCR was performed by using DreamTaq Green PCR master mix (Thermo Scientific, Waltham, MA, USA) according to the manufacturer’s protocol for 40 cycles using 10 mM primers and 5 µl sample. Primer sequences were as follows: ACGGGCGCGACCATCTGCGCTGC (forward, *cdt-1*), GCGAAGTACAGGTGGATGCTCTC (reverse, *cdt-1*), GCTGCTGCCGTCGGCGATCTTGC (forward, *gh1-1*), and ACGGGCGCGACCATCTGCGCTGC (reverse, *gh1-1*) (IDT, Coralville, IA, USA). PCR product and GeneRuler 1 kb DNA ladder (Thermo Scientific, Waltham, MA, USA) were run on a 1% agarose gel (Invitrogen, Thermo Scientific, Waltham, MA, USA) at 120V for 20 minutes and bands were visualized with SYBR safe DNA gel stain (Invitrogen, Waltham, MA, USA) on an iBright FL1500 imager (Invitrogen, Thermo Scientific, Waltham, MA, USA).

### Cuprizone Intoxication

Mice aged 8-10 weeks old were fed cuprizone diet for 6 weeks (Supelco, Sigma Aldrich, St. Louis, MO, USA). Cuprizone chow was prepared fresh daily by mixing powdered standard chow with 0.25% cuprizone (w/w). After 6 weeks of feeding, mice were sacrificed or returned to standard chow without cuprizone for 2-4 weeks.

### *Myo*-Inositol Supplementation of Animals

*Myo*-inositol supplementation was performed on animals following three weeks of 0.25% cuprizone supplementation. At the end of week three of cuprizone treatment, facility water was replaced with water containing 6 mg/liter, 600 mg/liter, or 6000 mg/L of *myo*-inositol (Thermo Scientific, Waltham, MA, USA). Animals were maintained on *myo*-inositol supplementation for weeks 4-6 of cuprizone treatment. Supplemented water was prepared fresh weekly. Control animals received facility water alone for the duration of the experiment.

### Primary Cell Culture

All cells were cultured at 37 °C and 5% CO_2_ in Poly-D-Lysine coated plates. Mouse primary cells were isolated from <P3 pups. In brief, brains were isolated and placed in Hibernate-A media supplemented with 0.5mM Glutamax and 2% B27 supplement (Gibco, Thermo Scientific, Waltham, MA, USA) and stored at 4 °C overnight. Tissue was dissociated into a single-cell suspension by using either Neural Tissue Dissociation Kit – Postnatal Neurons for isolation of neurons or Neural Tissue Dissociation Kit-P for OPC isolation (Miltenyi Biotec, Bergisch Gladbach, Germany). Magnetic bead separation was performed by using either the neuron isolation kit or CD140 microbead kit (Miltenyi Biotec, Bergisch Gladbach, Germany) for OPC isolation according to the manufacturer’s instructions. Neurons were cultured in standard neuron media consisting of MACS neuro media supplemented with 1X MACS NeuroBrew-21 (Miltenyi Biotec, Bergisch Gladbach, Germany) and 0.5 mM GlutaMAX (Gibco, Thermo Scientific, Waltham, MA, USA) unless otherwise stated. OPCs were cultured in standard neuron media supplemented with 10 ng/ml bFGF and 10 ng/ml PDGF (Miltenyi Biotec, Bergisch Gladbach, Germany) (standard OPC medium). Primary cell media were changed by half every other day, unless otherwise stated.

Neurons were cultured for 6 days in standard neuron media in a 24-well plate at 50,000 cells/well, with some wells containing no cells as controls for media evaporation. After 6 days, media was completely removed and replaced with Neurobasal-A media (without glucose, sodium pyruvate) supplemented with 1X B-27 Plus supplement, 0.5 mM GlutaMAX (Gibco, Thermo Scientific, Waltham MA, USA) and either 12.5 mM cellobiose (MP Biomedicals, Irvine, CA, USA), 25 mM glucose (Sigma Aldrich, St. Louis, MO, USA), or neither. Then, 10 µl media samples were collected at 4, 7, and 11 days, and media were replaced by half after collection on days 4 and 7. Viability was assessed after 7 days by using alamarBlue cell viability (Invitrogen, Thermo Scientific, Waltham, MA, USA) according to the manufacturer’s protocol and read with a Biotek Cytation 5 multimode reader (Agilent, Santa Clara, CA, USA).

### OPC Treatment

Primary OPCs were isolated from animals and plated at 200,000 cells/well in a 24-well plate and cultured for 5 days in standard OPC medium. For conditioned media experiments, cells were cultured in 1:1 neuron conditioned medium: standard OPC medium for up to 7 days. Control OPCs were cultured with non-conditioned neuron medium: standard OPC medium at a ratio of 1:1.

For *myo*-inositol treatment, OPCs were plated at 20,000 cells/well with additional wells containing no cells (growth medium only condition) in a 24 well plate and cultured for 7 days. Cells were subsequently treated with standard medium or medium containing 1 or 10 mM *myo*-inositol (Thermo Scientific, Waltham, MA, USA). After 7 days, cells were analyzed for immunofluorescence and proliferation was assessed by using a Cyquant direct cell proliferation assay (Invitrogen, Thermo Scientific, Waltham, MA, USA) under the same conditions. For RT-qPCR, cells were cultured at 200,000 cells/well and treated as above. After 7 days, cells were analyzed for RT-qPCR. For *myo*-inositol uptake, OPCs were plated at 50,000 cells/well in a 24-well plate. Next, 10 µl of medium was collected every 24 hours from wells cultured in 10 mM *myo*-inositol to measure uptake.

### RNAi Knockdown of SLC5A3 Expression

Oligodendrocyte precursor cells were plated at 200,000 cells/well in 24-well plates and cultured for 7 days in standard growth medium. Cells were treated with 1.5 µl lipofectamine RNAiMAX (Invitrogen, Thermo Scientific, Waltham, MA, USA) and 5 pmol siRNA (IDT, Coralville, IA, USA) in Opti-MEM medium (Gibco, Thermo Scientific, Waltham, MA, USA) per well and cultured for 3 days. Media were then removed by half and replaced with *myo*-inositol supplemented growth media to a final concentration of 5 mM and cultured for 3 days. New RNAi solutions were also added to each well on day 3. After the addition of medium containing *myo*-inositol, 10 µl samples were collected daily to measure *myo*-inositol uptake.

### Real-Time Quantitative RT-PCR Analysis

RNA was extracted by using TRI reagent with Direct-zol microprep kits for corpus callosum and miniprep kits for cells and whole brain (Zymo Research, Irvine, CA, USA) according to the manufacturer’s protocol. Synthesis of cDNA was performed by using SuperScript III First-Strand Synthesis Supermix (Invitrogen, Thermo Scientific, Waltham, MA, USA) according to the manufacturer’s protocol. Real-time quantitative RT-PCR was performed by using PowerUP SYBR Green Master Mix for qPCR (Applied Biosystems, Thermo Scientific, Waltham, MA, USA) according to the manufacturer’s protocol with melt curve and loaded with a normalized amount of cDNA. RT-qPCR was performed on a StepOnePlus Real-Time PCR system (Applied Biosystems, Thermo Scientific, Waltham, MA, USA). Primer information is listed in the resources table (Origene, Rockville, MA, USA). RT-qPCR samples were normalized by cDNA input. Data are represented as –(Delta Ct) values relative to GAPDH normalized to the maximum value at 100 and the minimum value at 0. Values returned as “undetected” were set to -35 as -DeltaCt.

### Seahorse Respiration Assay

Primary neurons were isolated and plated in Seahorse XFp PDL Cell Culture Miniplates (Agilent, Santa Clara, CA, USA) at 200,000 cells/well in growth medium. Cells were cultured at 37 °C and 5% CO2 for 3 days with media swapped by half every other day. On the day of the assay, media were swapped for Seahorse XF DMEM media (Agilent, Santa Clara, CA, USA) supplemented with 1 mM glutamine (Agilent, Santa Clara, CA, USA), 10 mM glucose (Agilent, Santa Clara, CA, USA), or 5 mM cellobiose (MP Biomedicals, Irvine, CA, USA) and 1 mM pyruvate (Agilent, Santa Clara, CA, USA) and plates were moved into a non-CO_2_ incubator for 45 minutes. Basal respiration was measured by using a hydrated sensor cartridge on a Seahorse XFp (Agilent, Santa Clara, CA, USA) according to the manufacturer’s instructions. After analysis, cells were counted under a microscope for data normalization.

### Stable Isotope Tracing

Primary neuronal cells were isolated and cultured in standard media for 5-7 days, media was fully removed and replaced with Neurobasal-A media supplemented with 0.5 mM Glutamax, 2% B-27 (Gibco, Thermo Scientific, Waltham, MA, USA), and either 25 mM [^13^C_6_] glucose or 12.5 mM [^13^C_12_] cellobiose (Omicron Biochemicals, South Bend, IN, USA). Media samples were collected at various time points and flash frozen before extraction with 2:2:1 Acetonitrile:Methanol:Water (Fisher Chemical, Waltham, MA, USA). Cell samples were collected by aspirating the media, washing twice with ice-cold PBS (Gibco, Thermo Scientific, Waltham, MA, USA) and once with ice-cold water, followed by quenching with ice-cold methanol. Cells were scraped and transferred to a new tube where acetonitrile and water were added to a final ratio of 2:2:1 and stored at -80°C until extraction.

Stable isotope tracing in animals was performed as follows. Prior to surgery, mice were injected with buprenorphine SR at 1 mg/kg. Solutions containing either 400 mM [^13^C_12_] cellobiose or 800 mM [^13^C_6_] glucose in artificial cerebrospinal fluid (CSF) (2.16 g/L NaCl, .056 g/L KCl, .052 g/L CaCl2 · 2H2O, .041 g/L MgCl2 · 6H2O, .214 g/L Na2HPO4 · 7H2O, and .007 g/L NaH2PO4 · H2O) (Sigma Aldrich, St. Louis, MO, USA) were prepared and used to fill osmotic pumps (Alzet, Durect Corporation, Cupertino, CA, USA). Pumps were assembled and primed for 1 hour at 37 °C. Mice were placed under 5% isoflurane until movement stopped and maintained at 2% isoflurane until surgery was finished. Mice were placed on a heated pad in a stereotactic frame zeroed to bregma, and cannulas were implanted at -1.1 mm lateral, -.4 mm posterior, and 2.5 mm depth. Cannulas were secured to the skull using Loctite 454, attached to the osmotic pump tubing as described by the manufacturer’s instructions. Incisions were nylon-stitched, treated with antibiotic ointment, and mice were placed in a heated recovery cage.

Following infusion, mice were placed under 5% isoflurane until movement stopped. Serum was collected via cardiac puncture and mice were immediately sacrificed by cervical dislocation. Brains were harvested, embedded in 5% carboxymethylcellulose (Fisher Scientific, Waltham, MA, USA) on dry ice or flash frozen in liquid nitrogen, and stored at -80 °C until extraction. Serum was placed on ice for 60 minutes to allow clotting and centrifuged at 14,000 rpm for 10 minutes to separate serum. Serum was stored at -80 °C until extraction.

### Metabolite Extraction

All solvents used were LC/MS grade (Fisher Chemical, Waltham, MA, USA). Media samples were extracted by using a 9:1 ratio of 2:2:1 Acetonitrile:Methanol:Water (AMW):sample, vortexed for 30 seconds, and stored at -20 °C overnight. Samples were centrifuged at 14,000 rpm for 10 minutes, and supernatant was transferred to LC vials for analysis. Serum was extracted in the same manner as above.

Cell samples suspended in 2:2:1 AMW underwent three freeze-thaw cycles with 10 minutes of sonication after each cycle and were then stored at -20 °C overnight. Samples were centrifuged at 14,000 rpm for 10 minutes, and supernatant was transferred to a new tube and dried via SpeedVac. Pellets were resuspended in 100 mM NaOH (Sigma Aldrich, St. Louis, MO, USA), heated at 95 °C for 5 minutes then cooled to room temperature for four cycles, and analyzed by using BCA assay (Pierce, Thermo Scientific, Waltham, MA, USA). Dried supernatant was resuspended in a normalized volume of AMW according to protein concentration, vortexed for 30 seconds, sonicated for 10 minutes at room temperature, and stored at -20 °C overnight. Samples were then centrifuged at 14,000 rpm for 10 minutes and transferred to LC vials for analysis via LC/MS.

Whole brain tissue for z-score analysis of pool sizes was performed using flash frozen tissue. Tissue was ground via mortar and pestle and kept frozen using liquid nitrogen. Frozen ground tissue was weighed and extracted using 40µl AMW per mg of tissue and following the sample protocol of freeze-thaw, storage, and centrifugation as above. Supernatant was transferred directly to LC for storage at -80°C until analysis.

Microdissection of corpus callosum was performed as follows. Tissue was sliced at 100 µm thickness on a CM1860 cryostat microtome (Leica Biosystems, Nussloch, Germany), and tissue region was excised via scalpel and transferred into pre-cooled tubes for extraction. Tissue was extracted by using 500 µl AMW and following the same method as above for cells.

### Metabolomics

All solvents used were LC/MS grade. Metabolite profiling was performed by using a Vanquish Horizon UHPLC system coupled to an Orbitrap ID-X Tribrid mass spectrometer (Thermo Fisher, Waltham, MA, USA). Metabolites were separated with an iHILIC-P classic column (100 x 2.1mm, 5µm, HILICON AB, Umeå, Sweden) at a column temperature of 45 °C and sample temperature of 6 °C. The following gradient was applied at a flow rate of 250 µL/min: 0-1 min, 90% B at 250 μL/min; 1-12 min, 35% B; 12-12.5 min, 25% B; 12.5-14.5 min, 25% B; 14.5-16.5 min, 90% B; 16.5-20.5 min, 90% B at 400 μL/min; 20.5-22 min, 90% B. Solvent A consisted of 95% water in 5% acetonitrile with 20 mM ammonium bicarbonate, 0.1% ammonium hydroxide, and 2.5 μm medronic acid. Solvent B consisted of 95% acetonitrile in 5% water. Metabolites were analyzed in positive and negative mode by using polarity switching under the following conditions: 67-1000 *m/z* scan, 120,000 resolution, 3500 V positive voltage, 2800 V negative voltage, 35 sheath gas flow rate, 10 auxiliary gas flow rate, 1 sweep gas flow rate, 60% RF lens, 300 °C ion transfer tube, and 200 °C vaporizer temp.

### Lipidomics

Lipidomics was performed by using a 1290 Infinity II UHPLC system coupled to an Agilent 6546 Q-TOF mass spectrometer (Agilent, Santa Clara, CA). Lipids were separated with an ACQUITY Premier HSS T3 column (100 x 2.1mm, 1.8µm, Waters, Milford, MA) at a column temperature of 55 °C and a sample temperature of 6 °C. The chromatographic gradient was linear and as follows: 0 min, 15% B at 400 μL/min; 0-2.5 min, 50% B; 2.5-2.6 min, 57% B; 2.6-9.0 min, 70% B; 9.0-9.1 min; 93% B; 9.1-11 min, 96% B; 11-11.1 min, 100% B; 11.1-12 min, 100% B; 12-12.2 min, 15% B; 12.2-16 min, 15% B. Solvent A consisted of 5:3:2 water:acetonitrile:isopropanol with 10 mM ammonium formate and 5 µM medronic acid. Solvent B consisted of 1:9:90 water:acetonitrile:isopropanol with 10 mM ammonium formate. Lipids were analyzed in positive mode under the following conditions: 200-1700 *m/z* scan, 1 spectra/second acquisition rate, 320 °C gas temp, 8 l/min drying gas, 45 psi nebulizer, 350 °C sheath gas temp, 11 l/min sheath gas flow, 3500 V capillary voltage, 1000 V nozzle voltage, 175 V fragmentor, 65 V skimmer, 750 V Oct 1 RF Vpp.

The analysis of PI lipid head groups was performed by using a Vanquish Horizon UHPLC system coupled to an Orbitrap ID-X Tribrid mass spectrometer (Thermo Fisher, Waltham, MA, USA). PI lipids were separated on an ACQUITY Premier HSS T3 column (100 x 2.1mm, 1.8 µm, Waters) at a column temperature of 45 °C and a sample temperature of 6 °C. The injection volume per sample was 10 µl. The linear chromatographic gradient was as follows: 0 min, 15% B at 400 μL/min; 0-2.5 min, 50% B; 2.5-2. 6 min, 57% B; 2.6-9.0 min, 70% B; 9.0-9.1 min, 93% B; 9.1-11 min, 96% B; 11-11.1 min, 100% B; 11.1-12 min, 100% B; 12-12.2 min, 15% B, 12.2-16 min, 15% B. Solvent A consisted of 5:3:2 water:acetonitrile:isopropanol with 10 mM ammonium formate and 5 µM medronic acid. Solvent B consisted of 1:9:90 water:acetonitrile:isopropanol with 10 mM ammonium formate. Lipids were analyzed in negative mode by using both MS1 and MS2. MS1 scan settings were as follows: from 4 - 9.5 minutes: 750-950 *m/z* scan, 80 ms maximum injection time, 60,000 resolution, 3000 V voltage, 50 sheath gas flow rate, 10 auxiliary gas flow rate, 1 sweep gas flow rate, 60% RF lens, 325 °C ion transfer tube, and 350 °C vaporizer temp. MS2 scan settings were as follows: from 0-16 minutes: quadrupole isolation mode with 1.6 m/z window, HCD activation with 35% collision energy, 30,000 resolution, 1500 ms maximum injection time. MS2 scan mass list table included highest intensity PIs (PI 18:0/20:4, *m/z* = 885.5499; PI 16:0/20:4, *m/z* = 857.5186) and their M+6 counterparts (*m/z* = 891.5700 and 863.5387, respectively) with z of 1. Fragment *m/z* = 247.032 represents the M+6 inositol head group fragment. A 5 ppm window was used for the extracted ion chromatograms.

### MALDI-Mass Spectrometry Imaging

Tissue sections of 10 µm were mounted on ITO-coated slides (Delta Technologies, Limited, Loveland, CO, USA) and dried under vacuum for 20 minutes. Matrix was applied by using an HTX M5 matrix sprayer (HTX imagine, Chapel Hill, NC, USA). Optical images were obtained on an Epson Perfection V850 Pro (Los Alamitos, CA, USA), and mass spectrometry images were generated by using a TimsTOF Flex (Bruker Daltonics, Billerica, MA, USA). For small molecule analysis, N-(1-naphthyl) ethylenediamine dihydrochloride (NEDC) (Sigma Aldrich, St. Louis, MO, USA) was sprayed under the following conditions: 7 mg/ml in 70/30 methanol/water, 8 passes, 3 liter/min gas flow, 0.12 ml/min flow rate, 1200 mm/min velocity, 3 mm track spacing, 70 °C needle temperature, and 35 °C tray temperature. Mass spectrometry imaging was performed at 50 µm raster size by using a 50 µm laser size with beam scan and 200 shots at 10,000 frequency and 85% power. Scan was 60-650 *m/z*, 600 Vpp collision RF, 10 eV collision energy, 5 eV ion energy, 5 µs pre-pulse storage, and 45 µs transfer time.

For lipid analysis, 9-aminoacridine (9-AA) (Sigma Aldrich, St. Louis, MO, USA) was sprayed under the following conditions: 5 mg/ml in 90/10 methanol/water, 8 passes, 3 liter/min gas flow, 0.11 ml/min flow rate, 700 mm/min velocity, 2 mm track spacing, 85 °C needle temperature, and 35 °C tray temperature. Imaging was performed at 30 µm raster size by using a 20 µm laser size with beam scan and 200 shots at 10,000 frequency and 50% power. Scan was 250-1150 *m/z*, 1800 Vpp collision RF, 10 eV collision energy, 7 eV ion energy, 7.0 µs pre-pulse storage, and 75 µs transfer time. Data were analyzed by using SCiLS Lab (Bruker Daltonics, Billerica, MA, USA) to generate .imzml files and a custom script to recalibrate pixels.

### Immunofluorescence

Cells were cultured on Poly-D-Lysine coated coverslips as described in the methods for primary cell culture. Cells were washed with ice-cold PBS three times before fixation with fresh 4% paraformaldehyde (Sigma Aldrich, St. Louis, MO, 63112) for 10 minutes at room temperature. Samples were washed 3x with ice-cold PBS (Gibco, Thermo Scientific, Waltham, MA, USA), permeabilized with .1% Triton X-100 (Sigma Aldrich, St. Louis, MO, USA) in PBS for 10 minutes, and washed 3x with ice-cold PBS. Blocking buffer (5% Bovine Serum Albumin (BSA) (Miltenyi Biotec, Bergisch Gladbach, Germany) in PBS) was applied for 90 minutes at room temperature. Primary antibodies were prepared in blocking buffer at the following concentrations: Alexa Fluor® 488 anti-GFAP Antibody (1:200), Alexa Fluor® 488 anti-Vimentin Antibody (1:100), and Alexa Fluor® 647 anti-Myelin Basic Protein Antibody (1:100), Alexa Fluor® 594 Anti-NeuN antibody (1:100), PE/Dazzle™ 594 anti-mouse CD140a Antibody (1:100), and Alexa Fluor® 647 anti-Tubulin β 3 (TUBB3) Antibody (1:200), followed by incubation overnight at 4°C. Coverslips were washed 3x with blocking buffer before being mounted on to Superfrost Plus slides (Fisherbrand, Thermo Fisher, Waltham, MA, USA) using Fluoromount G with DAPI or ProLong Glass Antifade Mountant with NucBlue Stain (Invitrogen, Thermo Scientific, Waltham, MA, USA). Slides were cured in the dark overnight before being sealed and imaged by using a DMi8 Thunder Imager and LAS X software (Leica Biosystems, Nussloch, Germany).

Tissue sections were sliced at a thickness of 10 µm by using a CM1860 cryostat microtome (Leica Biosystems, Nussloch, Germany) and thaw-mounted onto Superfrost Plus slides (Fisherbrand, Thermo Fisher, Waltham, MA, USA). Slides were placed in a humidity chamber at room temperature and fixed with fresh 4% paraformaldehyde (Sigma Aldrich, St. Louis, MO, 63112) for 10 minutes, washed 3x with ice-cold PBS (Gibco, Thermo Scientific, Waltham, MA, USA), permeabilized with .1% Triton X-100 (Sigma Aldrich, St. Louis, MO, USA) in PBS for 10 minutes, washed 3x with ice-cold PBS, and blocked with blocking buffer (5% Bovine Serum Albumin (BSA) (Miltenyi Biotec, Bergisch Gladbach, Germany) in PBS for 90 minutes. Primary antibodies were incubated overnight at 4 °C at the following dilution ratios of antibody to blocking buffer: Alexa Fluor® 488 anti-GFAP Antibody (1:200), Alexa Fluor® 647 anti-Neurofilament H & M (1:100), Alexa Fluor® 488 anti-Vimentin Antibody (1:100), PE/Dazzle™ 594 anti-mouse CD140a Antibody (1:100), Spark YG™ 570 anti-Myelin Basic Protein Antibody (1:100), Alexa Fluor® 647 anti-Myelin Basic Protein Antibody (1:100), and Alexa Fluor® 647 anti-TUBB3 Antibody (1:200) (all Biolegend, San Diego, CA, USA). Slides were washed with blocking buffer three times, and coverslips (Fisherbrand, Waltham, MA, USA) were mounted by using ProLong Glass Antifade Mountant with NucBlue Stain (Invitrogen, Thermo Scientific, Waltham, MA, USA). Slides were imaged by using a DMi8 Thunder Imager and LAS X software (Leica Biosystems, Nussloch, Germany).

### Public snRNA-seq Analysis

To examine metabolic gene changes between remyelination and demyelination in oligodendrocyte precursor cells, we reanalyzed the snRNA-seq dataset generated by Hou et al. (*31*). The raw matrix of UMI counts were downloaded from GEO under accession number of GSE204770 and analyzed by using the Scanpy package. Pathway enrichment analysis was performed on differentially expressed genes (DEGs) in OPCs and neurons at both the demyelination and remyelination time points. Genes with log2 (Fold Change) > 0.25 and adjusted p < 0.05 were considered as significant DEGs. Pathway analysis was performed on DEGs by using MetaboAnalyst (*59*) andiPath3 (*60*).

### Data Analysis

Metabolite identification and peak area extraction were performed by using Skyline software version 24.1 (*61*) and an in-house database to match retention time (±.5 min) and *m/z* (10 ppm). Untargeted analysis of isotope labeling was performed with X^13^CMS (*62*) For isotope tracing, natural abundance correction was performed by using AccuCor (*63*). Visualization was performed by using GraphPad Prism version 10.0 or Morpheus (https://software.broadinstitute.org/morpheus). Statistics were performed by using GraphPad Prism version 10.0. Statistical outliers were removed as determined via ROUT at Q= .1%.

### Code availability

Source code for the generation of fractional labeling from mass spectrometry imaging can be found on GitHub (https://github.com/e-stan/imaging) and is available under the Zenodo identifier https://doi.org/10.5281/zenodo.777844376. Metabolomics datasets are available at the Metabolomics Workbench (https://www.metabolomicsworkbench.org/).

## Acknowledgements

The authors thank the many members of the Patti and Johnson labs who contributed to this project over the past fifteen years. We are particularly grateful for the support we received during the early stages of the project’s development from the Pew Scholars Program in the Biomedical Sciences and the National Institutes of Health award R21CA191097 and award R35ES028365.

## Author Contributions

Conceptualization: G.J.P., S.L.J., L.P.S.; Methodology: G.J.P., S.L.J., L.P.S. K.A.T., R.F.G.; Formal analysis: K.A.T., M.S., M.S.H. K.C., R.F.G., S.L.J., L.P.S., G.J.P; Investigation: C K.A.T., M.S., M.S.H. K.C.,; Resources: G.J.P., S.L.J.; Data Curation: K.A.T., M.S., M.S.H., K.C., S.L.J., L.P.S, G.J.P.; Writing - Original Draft: L.P.S., G.J.P.; Writing - Review & Editing: K.A.T., M.S., K.C., R.F.G., L.P.S., G.J.P; Visualization: K.A.T., M.S., M.S.H., K.C., R.F.G., S.L.J., L.P.S., G.J.P; Supervision: S.L.J., L.P.S., G.J.P.; Funding Acquisition: S.L.J, G.J.P

---

**Fig. S1.**
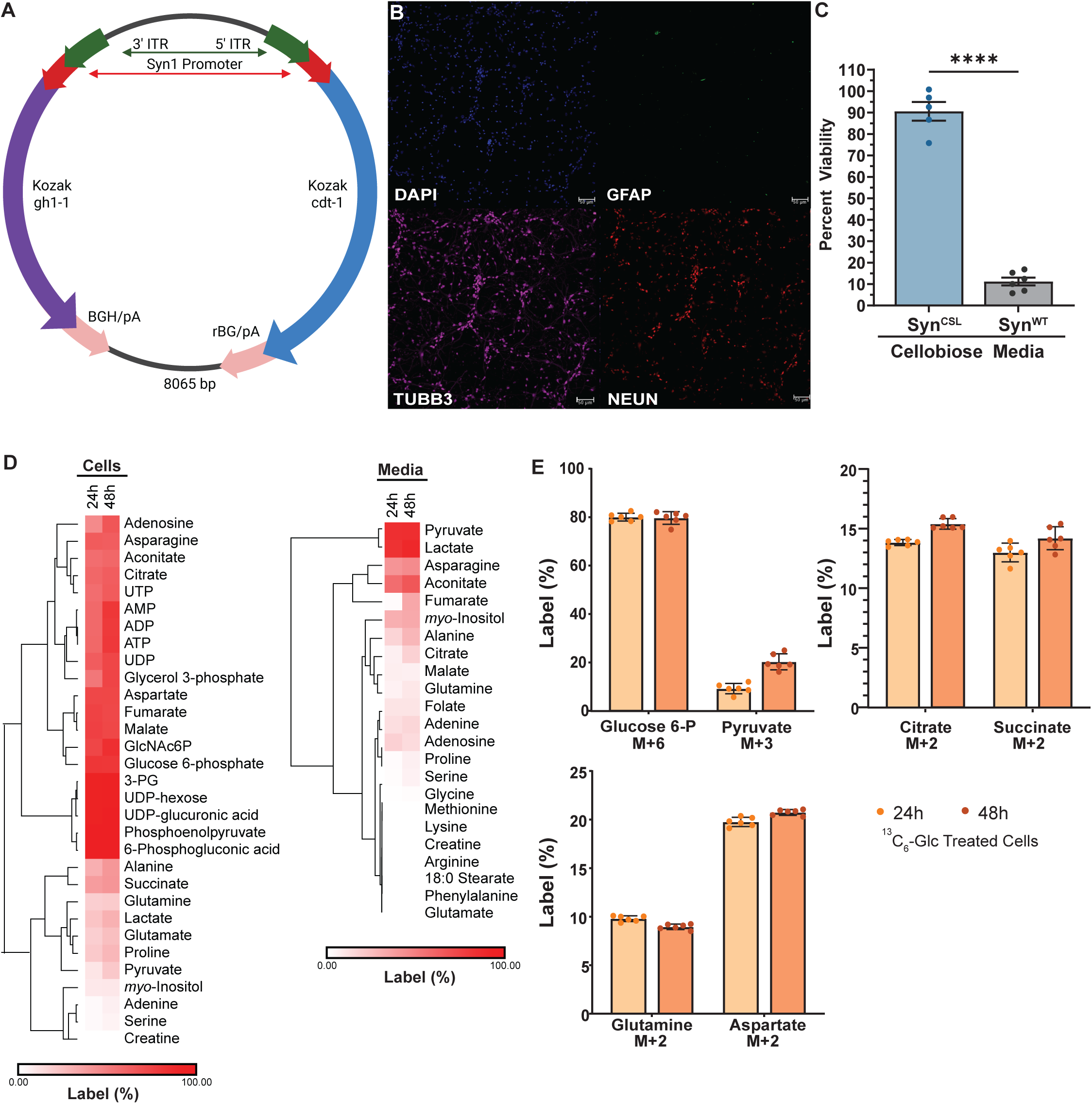
Validation of Syn^CSL^ transgenic neurons. **(A)** Diagram of the plasmid used to generate the Syn^CSL^ mouse line. **(B)** Representative immunofluorescence images of primary neuron culture for nuclei (DAPI-blue), astrocytes (GFAP-green), and neurons (TUBB3-pink, NEUN-red). Scale bars represent 50 µm, 1 million cells/well cultured for 7 days. **(C)** Viability measured by using the alamarBlue assay of Syn^CSL^ and wild-type neurons cultured in 12.5 mM cellobiose-containing media. Per well, 50,000 cells were plated and cultured for 7 days. Data normalized to viability of glucose-treated neurons (Syn^CSL^, n=5 wells; Syn^WT^, n=6 wells). Significance determined via two-tailed t-test. **(D)** Heatmap of average total percent labeling of metabolites in primary neurons (left) and media (right) when cultured for 24 or 48 hours with 25 mM ^13^C_6_ -glucose. Per well, 1 million cells were plated. Euclidean clustering of rows is based on average. **(E)** Percent labeling of M+6 G6P, M+3 pyruvate, M+2 citrate, M+2 succinate, M+2 glutamine, and M+2 aspartate in primary neurons cultured in 25 mM ^13^C_6_ -glucose for 24 or 48 hours. Per well, 1 million cells were plated (n=6 wells per group (D, E)). Percent labeling determined by LC/MS and corrected for natural abundance (D, E). Bars represent mean±SEM (C, E). (* p<.05, ** p<.01), *** p<.001, **** p<.0001).

**Fig. S2.**
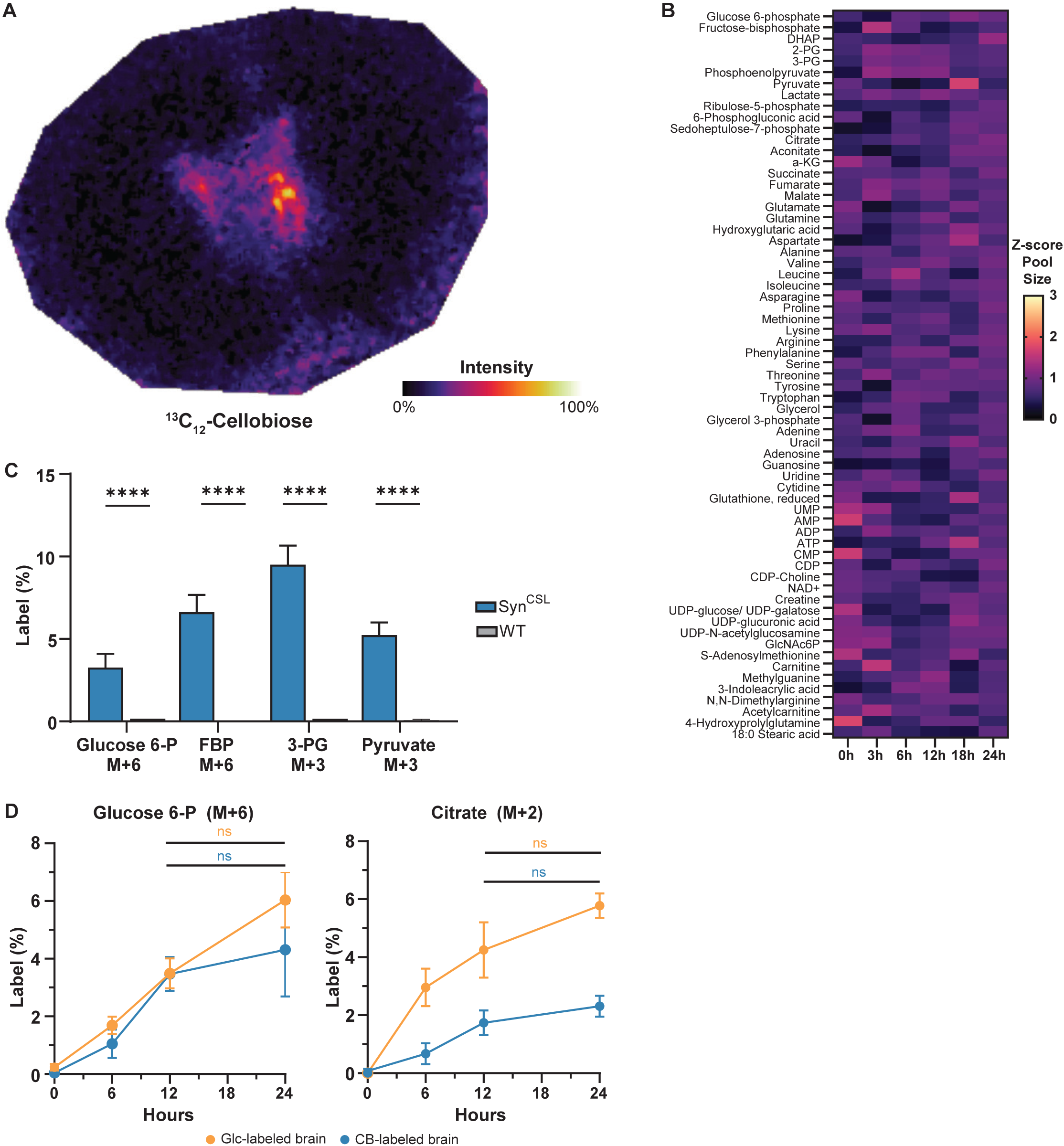
ICV tracer administration effectively tracks metabolism without perturbing metabolic homeostasis. **(A)** MALDI image of ^13^C_12_-cellobiose shows that the tracer is centered around lateral ventricles. Image shown is from the chloride adduct of cellobiose containing twelve ^13^C labels with weak denoising. It was generated from a Syn^CSL^ mouse administered 400 mM ^13^C_12_ -cellobiose at 8 µl/hour for 24 hours. **(B)** Heatmap of Z-scores of metabolite pools in brains of mice over the course of glucose administration relative to 0-hour control. (0h and 3h, n= 4 animals; 18h, n=5; 6h and 12h, n=6; 24h, n=11). **(C)** Percent of M+6 G6P labeling, M+6 FBP labeling, M+3 3-phospho-glycerate (3-PG) labeling, and M+3 pyruvate labeling from Syn^CSL^ and wild-type mice administered 400 mM ^13^C_12_ -cellobiose at 8µl/hour for 24 hours (n=5 animals per group). **(D)** Percent of M+6 G6P labeling and M+2 citrate labeling from brains after administration of 400 mM ^13^C_12_ -cellobiose or 800mM ^13^C_6_ -glucose. (CB 0h, n=6 mice; Glc 0h, n=15; CB 6h and Glc 12h, n=8; Glc 6h and CB 24h, n=10; CB 12h, n=9; Glc 24h, n=12). Significance of 12 vs. 24 hours determined via Mann Whitney test with Holm-Sidak correction (C, D). Significance of glucose vs. cellobiose determined via functional data analysis (C, D) (p-value G6P=.472; p-value citrate=.002). Data represents mean±SEM (C, D). (* p<.05, ** p<.01, *** p<.001, **** p<.0001).

**Fig. S3.**
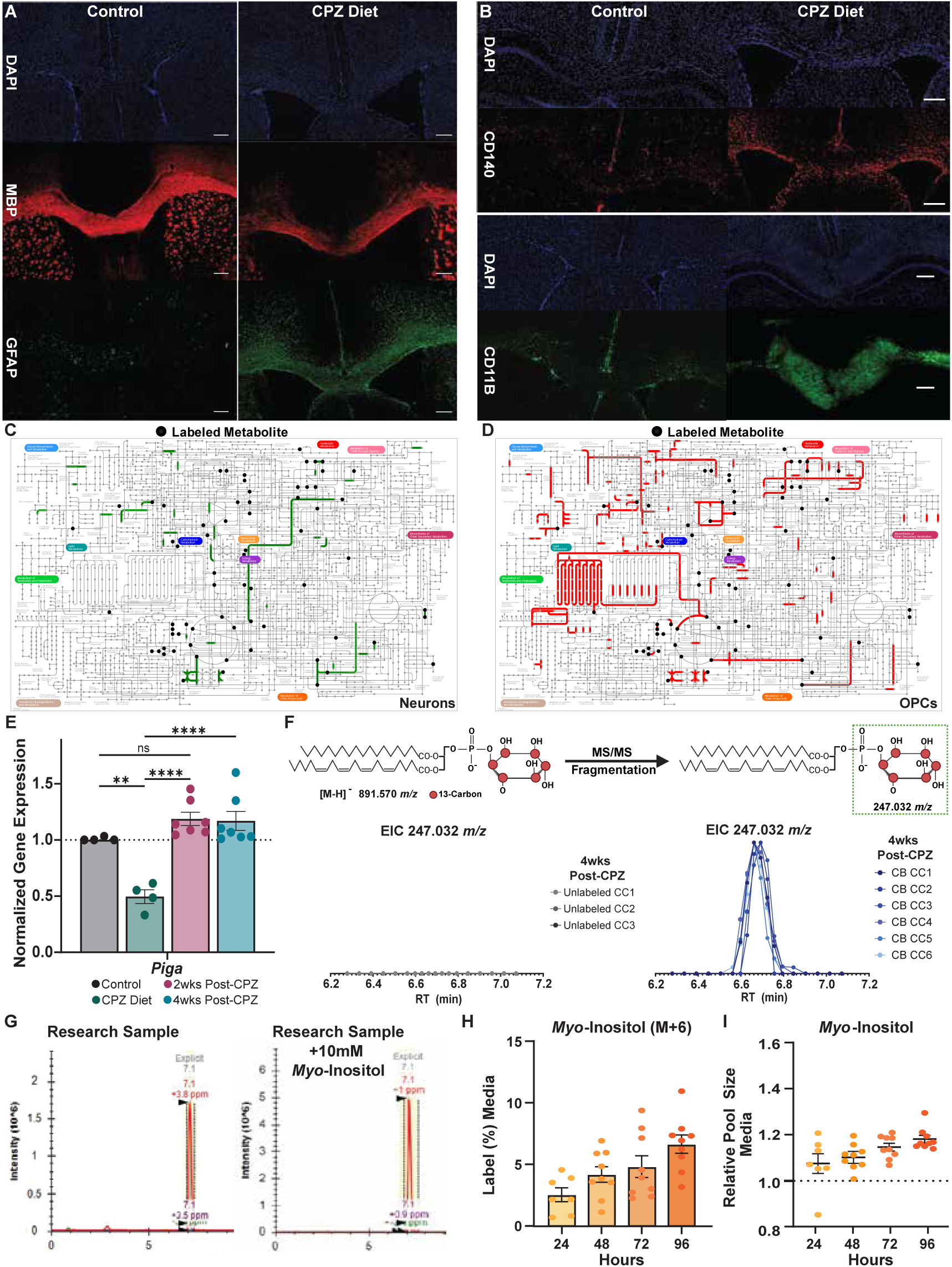
Demyelination and remyelination alter inositol metabolism. **(A)** Representative immunofluorescence images of brains from C57BL/6J mice fed standard or CPZ diet for 6 weeks and stained for nuclei (DAPI-blue), myelin (MBP-red), and astrocytes (GFAP-green). **(B)** Immunofluorescence images of brains from Syn^WT^ mice fed standard or CPZ diet for 6 weeks and stained for nuclei (DAPI-blue), OPCs (CD140-red), and microglia (CD11B-green). Scale bars represent 250 µm (A, B). **(C, D)** Integrative pathway mapping of DEGs and labeled metabolites in neurons (C) and OPCs (D). DEGs derived from snRNA-seq and labeled metabolites measured by LC/MS after ^13^C_12_-cellobiose delivery. Colored lines represent genes increased in expression (neurons-green, OPCs-red) during remyelination as compared to demyelination. Black circles indicate labeled metabolites during remyelination. Images generated with iPath3. **(E)** Normalized gene expression of *Piga* in the corpus callosum of mice fed standard diet (control), 6 weeks CPZ diet, or 6 weeks CPZ followed by 2 or 4 weeks standard diet (2 and 4wks post-CPZ, respectively). Data normalized to control mice fed standard diet by using the *Gapdh* housekeeping gene (control and CPZ diet, n=4 animals per groups; 2wks and 4wks post-CPZ, n=7). Significance determined via one-way ANOVA with Tukey’s correction. **(F)** Extracted ion chromatogram (EIC) of MS/MS fragment 247.032 *m/z* (±5 ppm), representing the M+6 inositol phosphate head group, in corpus callosum (CC) samples taken at 4-wks post-CPZ diet. Fragment derived from M+6 PI 18:0/20:4 (precursor *m/z* 891.570±10 ppm). Left EIC from unlabeled brains and right EIC from cellobiose-treated brains. **(G)** EIC of *m/z* =179.056 (±5ppm) in a QC sample made by pooling research specimens. EIC from the same sample spiked with 10 mM *myo*-inositol standard for confirmation of identification. **(H)** M+6 *myo*-inositol labeling from the media of primary neurons cultured in 25 mM ^13^C_6_ -glucose corrected against 0 h timepoint. **(I)** Relative *myo*-inositol pool size in culture media of primary neurons given 25 mM ^13^C_6_ -glucose normalized to 0 hour time-point. Per well, 1 million cells were plated (24 hours, n=7 wells; 48, 72, and 96 hours, n=9. (H, I)). Data are expressed as mean±SEM (E, H, I). (* p<.05, ** p<.01), *** p<.001, **** p<.0001).

**Fig. S4.**
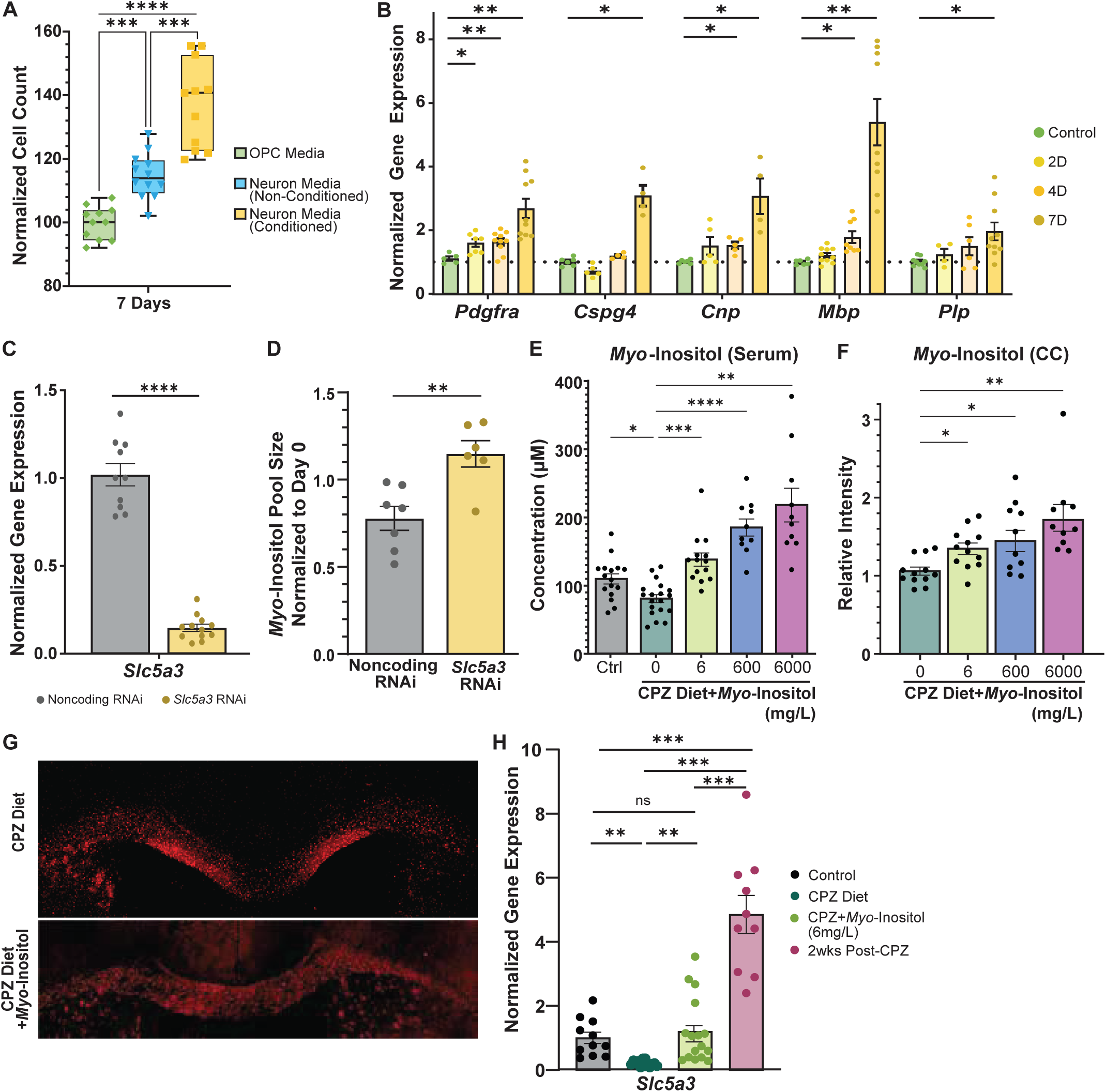
***Myo*-inositol transfer through SLC5A3 increases during remyelination. (A)** Normalized cell count of OPCs cultured in neuron-conditioned media for 7 days relative to unconditioned neuron media or OPC media (n=11 per condition). Error bars represent min to max. **(B)** Normalized gene expression of OPCs cultured in neuron-conditioned media for up to 7 days. Per well, 200,000 OPCs were cultured for 7 days in media conditioned by 1 million neurons/well for 7 days. (control, n≥5; 2D, n≥4; 4D and 7D, n≥4 cultures). Expression is normalized to control OPCs cultured in standard OPC media with the *Gapdh* housekeeping gene. Significance determined by using Welch’s t-test with Holm-Sidak correction. **(C)** Normalized gene expression of *Slc5a3* in OPCs (200,000 cells/well) treated with noncoding RNAi or *Slc5a3* RNAi for 6 days. Data normalized to noncoding RNAi with the *Gapdh* housekeeping gene (noncoding RNAi, n=10; *Slc5a3* RNAi, n=12). Significance determined via Student’s t-test. **(D)** Normalized pool size of *myo*-inositol in the media of OPCs treated with noncoding or *Slc5a3* RNAi for three days. Pool size normalized to 0-day timepoint; samples taken on day 3 after addition of *myo*-inositol supplemented (5mM) media. Significance determined via Welch’s two-tailed t-test (noncoding RNAi, n=6 and *Slc5a3* RNAi, n=7). **(E)** Concentration of *myo*-inositol measured by LC/MS in the serum of mice receiving standard chow (control) or CPZ diet with *myo*-inositol supplemented water (0, 6, 600, or 6000 mg/L *myo*-inositol). (control, n=16; CPZ, n=19 mice; 6mg/L *myo*-inositol, n=14; 600 mg/L and 6000 mg/L *myo*-inositol, n=10 mice). **(F)** Relative intensity measured by LC/MS of *myo*-inositol in the corpus callosum of mice fed a 6-week CPZ diet with 0, 6, 600 or 6000 mg/L *myo*-inositol supplemented water co-administered for the final three weeks of diet. Intensity relative to corpus callosum of CPZ-fed mice (0 mg/L and 6 mg/L, n=12 mice; 600 mg/L and 6000 mg/L, n=10 mice). **(G)** Representative immunofluorescence images of myelin (MBP-red) in brains of Syn^WT^ mice fed a 6-week CPZ diet with or without *myo*-inositol supplementation (6000 mg/L) for the final 3 weeks. Scale bars represent 500 µM. **(H)** Normalized gene expression of *Slc5a3* in corpus callosum of mice that received standard diet (control), 6 weeks of CPZ diet, CPZ diet with co-supplementation of 6 mg/L *myo*-inositol for the final 3 weeks (CPZ+*myo*-inositol), or an additional 2 weeks of standard diet after CPZ diet (2wks post-CPZ). Data normalized to control mice fed standard diet by using the *Gapdh* housekeeping gene (control, n=11 mice; CPZ diet, n=14; CPZ+*myo*-inositol, n=16; 2wks post-CPZ, n=10 animals). Bars represent mean±SEM (B, C, D, E, F, H). Significance determined using Brown-Forsythe and Welch ANOVA with Games-Howell’s post hoc (A, H) or with Dunnett’s correction (B, E, F). (* p<.05, ** p<.01), *** p<.001, **** p<.0001).

